# Multi-timescale Computation by Astrocytes

**DOI:** 10.64898/2026.01.05.697758

**Authors:** Chang Li, Lulu Gong, Chenghui Song, ShiNung Ching, Lucas Pozzo-Miller, Wei Li

## Abstract

The basic computational unit of the brain has long been defined as the neuron. However, mounting evidence suggests that other cells, especially astrocytes, also perform computations. Here, we demonstrate that cerebellar astrocytes decompose norepinephrine input into slow and fast calcium ion activities through differential adrenergic receptor engagement. During reward learning in mice, these slow and fast activities selectively target and modulate distinct synaptic pathways. Causal manipulations reveal that fast α2-adrenergic signals govern event-triggered responses and reinforcement learning, whereas slow α1-adrenergic signals maintain behavioral states and coordinate transitions. Remarkably, an actor-critic neural network trained on a similar sequence task spontaneously recapitulates these multitemporal dynamics, suggesting that astrocytes implement critic-like computations that evaluate states and modulate neuronal learning. These findings establish astrocytes as multilevel processors that transform univariate neuromodulatory inputs into multivariate, pathway-specific circuit controls operating in parallel with neuronal processing.

## Main

Neurons have traditionally been considered the fundamental computational units of the brain^1–3^. However, astrocytes exhibit signaling spanning orders of magnitude in time^4,5^, contact thousands of synapses^6–8^, and receive neuromodulatory inputs through diverse receptors^9,10^, suggesting that they may also perform computations.

Traditional characterizations of astrocytic calcium ion (Ca²⁺) signals as slow, uniform events^11–13^ have supported their modulatory roles in adjusting circuit tone^14,15^. Recent studies have identified faster transients^16–18^, but whether these signals transform neuromodulatory inputs into functionally distinct outputs that causally regulate behavior remains uncharacterized^19–21^. The cerebellar cortex provides an ideal model: climbing fibers (CFs) carry instructive signals^22,23,^ and parallel fibers (PFs) convey contextual information^24,25^, converging onto and making synaptic contacts with Purkinje cells, the sole output of the cerebellar cortex^26,27^. Bergmann glia (BG), the neighboring astroglia of Purkinje cells, interact with both synapses^28,29^, positioning them to differentially regulate these pathways.

Here, we show BG decompose norepinephrine signaling into slow and fast calcium ion activities through different α-adrenergic receptors, with each selectively modulating distinct synaptic pathways and controlling separate behavioral functions. Causal manipulations confirm that disrupting each activity mode impairs distinct aspects of behavior. An actor-critic network trained on a similar task spontaneously develops these multi-timescale dynamics, suggesting that astrocytes implement computations that operate in parallel with neurons to coordinate behavior.

### Cerebellar Astrocyte Ca²⁺ Dynamics During Reward-Guided Behavior

To examine Bergmann glia (BG) Ca²⁺ activity during behavior, we trained mice on a spatial sequence task requiring movement from a trigger zone to a reward zone (Fig. 1a). Sequence completion triggered sucrose delivery after 2 s, followed by a 4 s reward window. The mice were trained for 30 min per day for 15 days, with Day 1 serving as the no-reward baseline. Infrared sensors detected trigger and reward zone entry/exit, reward delivery, and licking, enabling precise alignment of behavior and BG Ca²⁺ activities.

**Figure 1:**
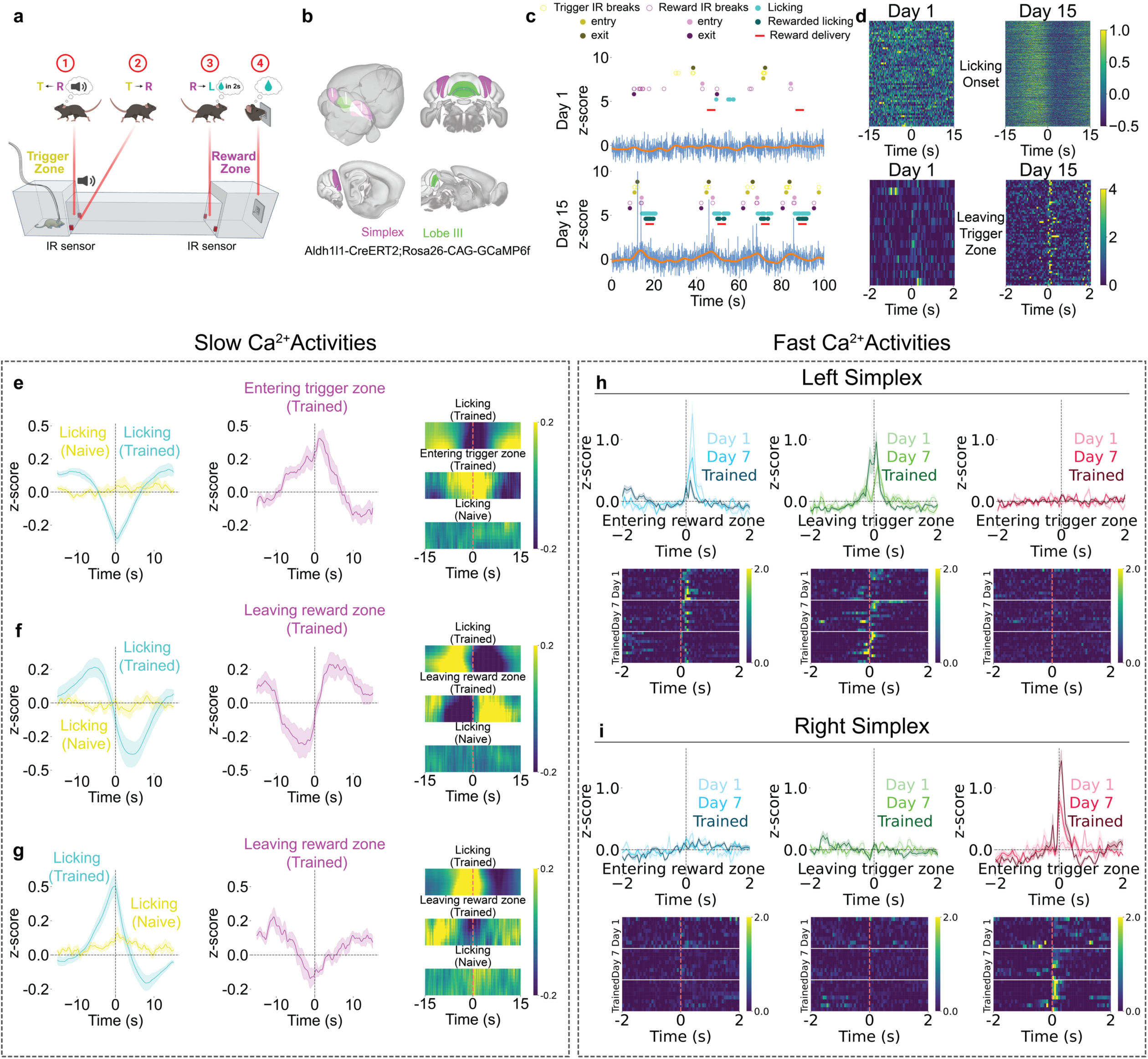
Cerebellar Astrocyte Ca²⁺ Dynamics During Reward-Guided Behavior. **(a)** Schematic of the spatial sequence task showing four trial phases: entering the trigger zone with auditory tone, leaving the trigger zone toward the reward zone, entering the reward zone with a 2-second delay, and licking during the 4-second reward window. IR sensors monitor zone transitions and licking (BioRender). **(b)** Fiber photometry recording sites in lobule III (green, medial vermis) and simplex (purple, lateral hemispheres) of *Aldh1l1*::CreERT2 x *Rosa26*::CAG-GCaMP6f mice expressing GCaMP6f in Bergmann glia (HERBS, Allen Mouse Brain Atlas, and BioRender). **(c)** Representative Ca²⁺ traces (z-scored ΔF/F) from Days 1 and 15. Blue: raw fluorescence. Orange: low pass filtered (0.3 Hz) slow oscillations. Behavioral markers: reward zone IR breaks (purple), trigger zone IR breaks (yellow), licking (cyan), and reward delivery (red). For zone IR breaks: hollow circles represent all breaks; light solid circles represent first break (entering); and dark solid circles represent last break (leaving). Day 15 shows robust task-aligned signals absent on Day 1. **(d)** Heatmaps of activity (z-scored ΔF/F) aligned to licking onset (top) and leaving the trigger zone (bottom), showing the emergence of slow ramping signals and fast transients with training. **(e-g)** Three slow signal types (z-scored ΔF/F; curve plots and heatmaps). **(e)** Trough signals (n=8 sites, 7 animals) decline before licking and peak at trigger zone entry. **(f)** State-change signals (n=8 sites, 6 animals) reverse polarity across reward transitions. **(g)** Peak signals (n=8 sites, 5 animals) ramp upward before licking and decline toward the reward zone exit. **(h,i)** Fast Ca²⁺ transients (z-scored ΔF/F) show hemisphere-specific training-dependent changes. Left simplex (n=11 sites, 11 animals): **(blue)** Entering the reward zone response diminishes with training. **(green)** Leaving the trigger zone response emerges with training. **(red)** Entering the trigger zone/tone onset results in a minimal response. Right simplex (n=8 sites, 8 animals): **(blue)** Entering the reward zone shows minimal modulation. **(green)** Leaving the trigger zone shows no significant response. **(red)** Entering the trigger zone/tone onset response increases with training.

We recorded BG Ca²⁺ via fiber photometry^30,31^ in *Aldh1l1*::CreERT2 x *Rosa26*::CAG-GCaMP6f induced by tamoxiphen^32,33^ recording from left and right simplex and lobule III (Fig. 1b; Extended Data Fig. 1). The animals progressively completed more trials, returned faster to the trigger zone, and restricted licking to the reward window, demonstrating the emergence of goal-directed behavior (Extended Data Fig. 2).

Example recordings show progression across training: Day 1 exhibited variable behavior with minimal signals, whereas Day 15 showed robust, temporally structured behavior and Ca²⁺ activity (Fig. 1c). The raw traces contained both slow ramping and brief fast transients; the filtered traces highlighted slower dynamics. Heatmaps aligned to the licking onset and trigger zone exit (Fig. 1d) show that slow ramping emerged during licking with training, whereas fast transients became prominent at the trigger zone exit.

### Slow BG Ca²⁺ Activities Encode Behavioral State Transitions

We classified slow BG Ca²⁺ dynamics during licking into three types—trough, state-change and peak—based on responses at the lick onset in trained animals (Fig. 1e-g). Trough signals declined before licking, reached a minimum at onset, and peaked at the trigger zone entry, suggesting reward suppression. State-change signals transitioned from positive to negative across the lick window and reversed polarity at the reward zone exit, which is consistent with shifts between reward seeking and engagement. Peak signals increased before licking, peaked at onset, and then declined toward the reward zone exit, which was consistent with reward activation. All three signal types appeared across cerebellar regions without clear anatomical bias (Extended Data Fig. 3). Notably, summing the average peak and trough traces produced a composite resembling the state-change trace, suggesting that state-change dynamics arise from the integration of opposing responses across regions.

Re-aligning traces to reward onset and performing slope analysis (Extended Data Fig. 4a) revealed consistent temporal trends across the pre-reward (−4 to 0 s), reward (0 to 4 s), and post-reward (4 to 8 s) periods. Trough signals were modulated before and after reward, whereas peak and state-change signals shifted during and after delivery. A comparison of rewarded and unrewarded spontaneous licking revealed that reward presence strongly influenced dynamics, with differences emerging before reward onset, suggesting preparatory components (Extended Data Fig. 4b-d).

### Fast BG Ca²⁺ Transients Show Hemisphere-Specific Tuning

Fast BG Ca²⁺ transients exhibited spatial distribution and learning modulation. In the left simplex lobule, reward zone entry evoked strong responses on Day 1 that diminished with training (Fig. 1l). Conversely, the trigger zone exit—the preceding reward approach—elicited signals that increased with training (Fig. 1m). No response occurred at trigger zone entry (tone onset), suggesting that auditory cues alone did not drive BG activity (Fig. 1n).

In the right simplex lobule, fast signals aligned to trigger zone entry (tone onset) and gradually increased with training (Fig. 1q), whereas responses to reward zone entry and trigger zone exit remained flat (Fig. 1o-p). The raw traces revealed prominent fast events in hemispheres but not in the midline lobule III (Extended Data Fig. 5).

### NE-Dependent Modulation of BG Ca²⁺ Activities

We tested whether norepinephrine (NE) modulates these activities during behavior. NE is a key neuromodulator released from locus coeruleus projections to the cerebellum^34–36^ and is implicated in cerebellar learning^37–39^. Using GRAB_NE sensors^40^ (Fig. 2a), we found that NE transients followed the same timing and trends as the three slow BG signal types did (Fig. 2b-d), indicating that slow BG Ca²⁺ tracks NE fluctuations. Re-alignment to reward onset revealed that NE displayed similar patterns but with sharper changes during reward (Fig. 2e).

**Figure 2:**
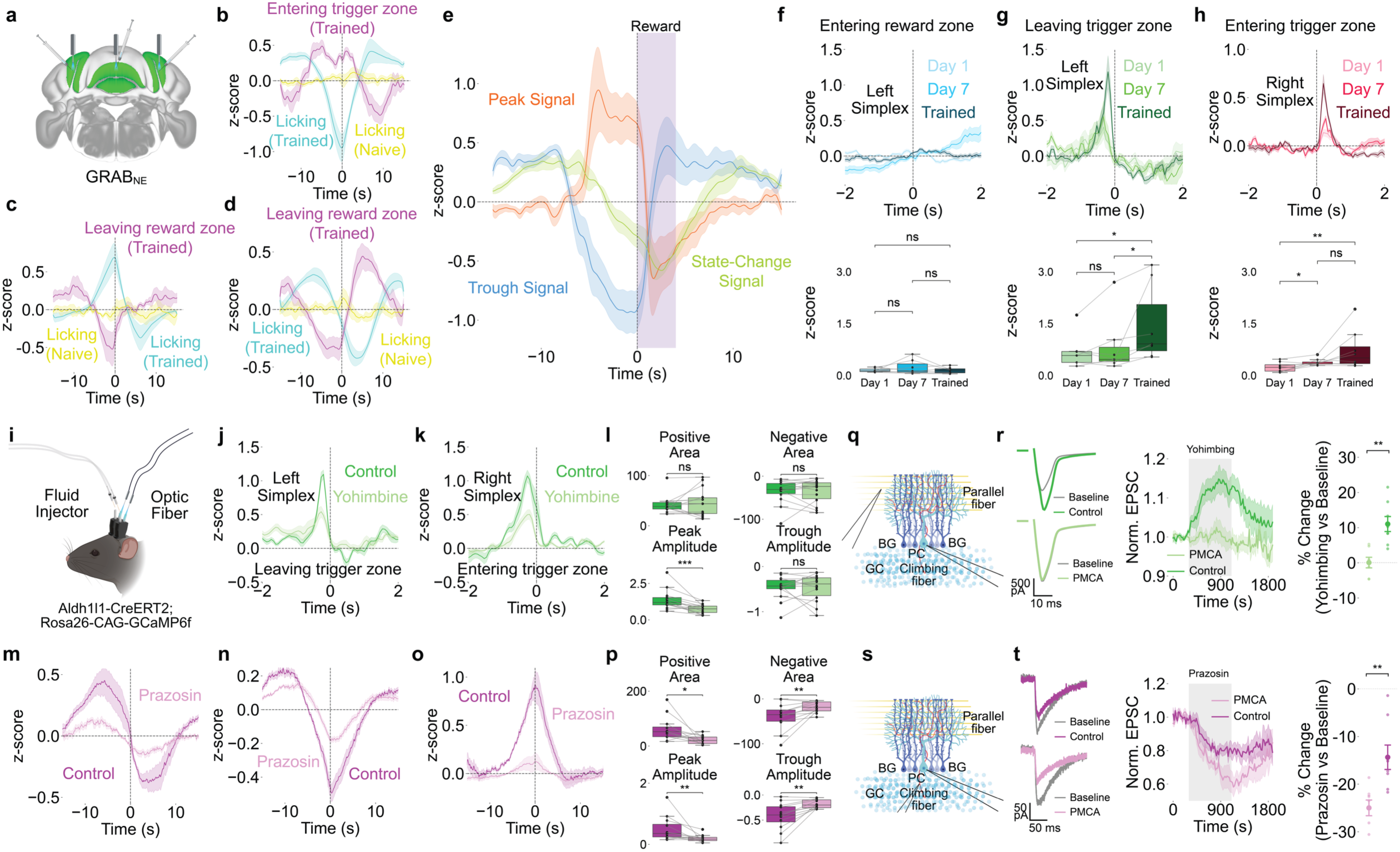
Norepinephrine-Dependent Modulation Through Distinct Adrenergic Receptors. **(a)** Schematic of GRAB_NE sensor expression in simplex and lobule III for measuring NE dynamics (HERBS, Allen Mouse Brain Atlas, and BioRender). **(b-d)** Slow NE dynamics (z-scored ΔF/F) show three signal types: **(b)** trough (n=11 sites, 7 animals), **(c)** peak (n=7 sites, 6 animals), and **(d)** state-change (n=6 sites, 5 animals) patterns. **(e)** Slow NE signals (z-scored ΔF/F) aligned to reward onset showing trough, peak, and state-change temporal profiles. **(f-h)** Fast NE dynamics (z-scored ΔF/F) showing hemisphere-specific training-dependent changes (n=8 sites per hemisphere). Left hemisphere: **(f)** minimal reward zone entry response, **(g)** emerging trigger zone exit response with training. **(h)** The right hemisphere shows progressive tone onset increases. **(i)** Combined fiber photometry and microinfusion system schematic (BioRender). **(j-l)** Yohimbine (an α2-AR antagonist) suppresses fast Ca²⁺ transients (z-scored ΔF/F) in **(j)** the recordings from the left hemisphere leaving trigger zone and **(k)** the recordings from the right hemisphere as the mice entering the trigger zone (n=14 sites, 8 animals). **(m-p)** Prazosin (an α1-AR antagonist) disrupts slow Ca²⁺ signals (z-scored ΔF/F): **(m)** state-change, **(n)** trough, and **(o)** peak signals with reduced amplitudes (n=12 sites, 7 animals). **(q-r)** Ex vivo climbing fiber EPSCs (n=8 control, n=7 PMCA cells, 5 animals each). Yohimbine enhances CF transmission, whereas astrocytic PMCA reduces it. **(s-t)** Ex vivo parallel fiber EPSCs (n=8 cells, 5 animals each). Prazosin and astrocytic PMCA reduce PF transmission.

Fast NE dynamics aligned to trigger zone exit (left hemisphere, Fig. 2g) and trigger zone entry (right hemisphere, Fig. 2h) and showed training-dependent activity similar to that of astrocytic fast transients. Reward zone entry showed a minimal NE response (Fig. 2f), which differed from the Ca²⁺ response. These findings suggest that NE acts as a reinforcement signal during cue presentation and the reward approach.

### α1-ARs and α2-ARs Mediate Distinct BG Ca²⁺ Dynamics

We combined fiber photometry with local drug infusion to test adrenergic receptor contributions (Fig. 2i). Yohimbine (an α2-AR antagonist) decreased the fast BG Ca²⁺ amplitude at the trigger zone exit (left hemisphere) and entry (right hemisphere, Fig. 2j-l), confirming that α2-ARs mediate fast transients. Yohimbine did not affect slow activity (Extended Data Fig. 6).

Prazosin (an α1-AR antagonist) decreased slow signals across events (Fig. 2m-p), reducing peak amplitude and shifting polarity balance, confirming that α1-ARs mediate slow dynamics. Prazosin did not affect fast transients (Extended Data Fig. 6). Propranolol (a β-AR antagonist) affected neither fast nor slow activity (Extended Data Fig. 6).

### Pathway-Specific Effects on Purkinje Cell Synaptic Inputs

Ex vivo patch-clamp recordings provided pathway-specific validation. For climbing fiber (CF) stimulation (Fig. 2q-r), yohimbine increased CF-evoked excitatory postsynaptic currents (EPSCs) while expressing plasma membrane Ca²⁺ ATPase (PMCA)^41,42^ in the BG via adeno-associated virus (AAV) injection—which enhances Ca²⁺ export from astrocytes and reduces astrocytic Ca²⁺ activity—significantly reduced responses.

For parallel fiber (PF) stimulation (Fig. 2s-t), prazosin decreased PF-evoked EPSCs, and PMCA expression further reduced them.

Immunohistochemistry confirmed that α2-ARs were highly expressed in the BG, α1-ARs co-labeled with S100β in astrocytes, and α1-ARs were adjacent to PF terminals marked by VGLUT1 (Extended Data Fig. 7). Conditional α1-AR knockout in the BG nearly eliminated α1-AR-evoked Ca²⁺ activity in the cerebellar cortex^43^, demonstrating functional localization to astrocytes adjacent to PF synapses. Prazosin did not affect CF-evoked EPSCs, which is consistent with α1-AR localization to PF pathways^23,24^, whereas yohimbine significantly reduced PF-evoked EPSCs during PF stimulation (Extended Data Fig. 6). Given that CF signals are classically viewed as teaching signals^44–46^ because they trigger complex spikes that suppress simple spikes from PF input, the observation that yohimbine enhances CF input while simultaneously suppressing PF input suggests that α2-AR signaling coordinates the balance between these pathways.

The temporal properties align with BG dynamics: granule cells providing PF input generate sparse, sustained contextual signals^47^ matching slow BG activities, whereas CF inputs deliver sharp, temporally precise instructive signals^47^ matching fast transients. These findings suggest that slow α1-AR-mediated signals regulate contextual PF pathways, whereas fast α2-AR-mediated signals regulate instructive CF pathways.

### Fast and Slow BG Activities Differentially Regulate Reward-Guided Behavior

We selectively perturbed fast α2-AR-dependent or slow α1-AR-dependent dynamics during the reward-guided task. For fast activity disruption, we expressed ChETA-EYFP^48,49^ in the BG and delivered optogenetic stimulation upon sensor breaks in trigger or reward zones during Days 1-10, followed by no-stimulation recovery (Days 11--15; Fig. 3a-c). This produced specific impairments: ChETA animals presented more trigger zone crossings, more sensor breaks, less reward licking, a lower rewarded-to-total lick ratio, and increased trigger-to-reward time, which was consistent with impaired reinforcement learning and altered reward behavior. Impairments partially reversed when stimulation ceased, indicating that fast BG activities transiently shape task execution rather than causing long-lasting deficits.

**Figure 3:**
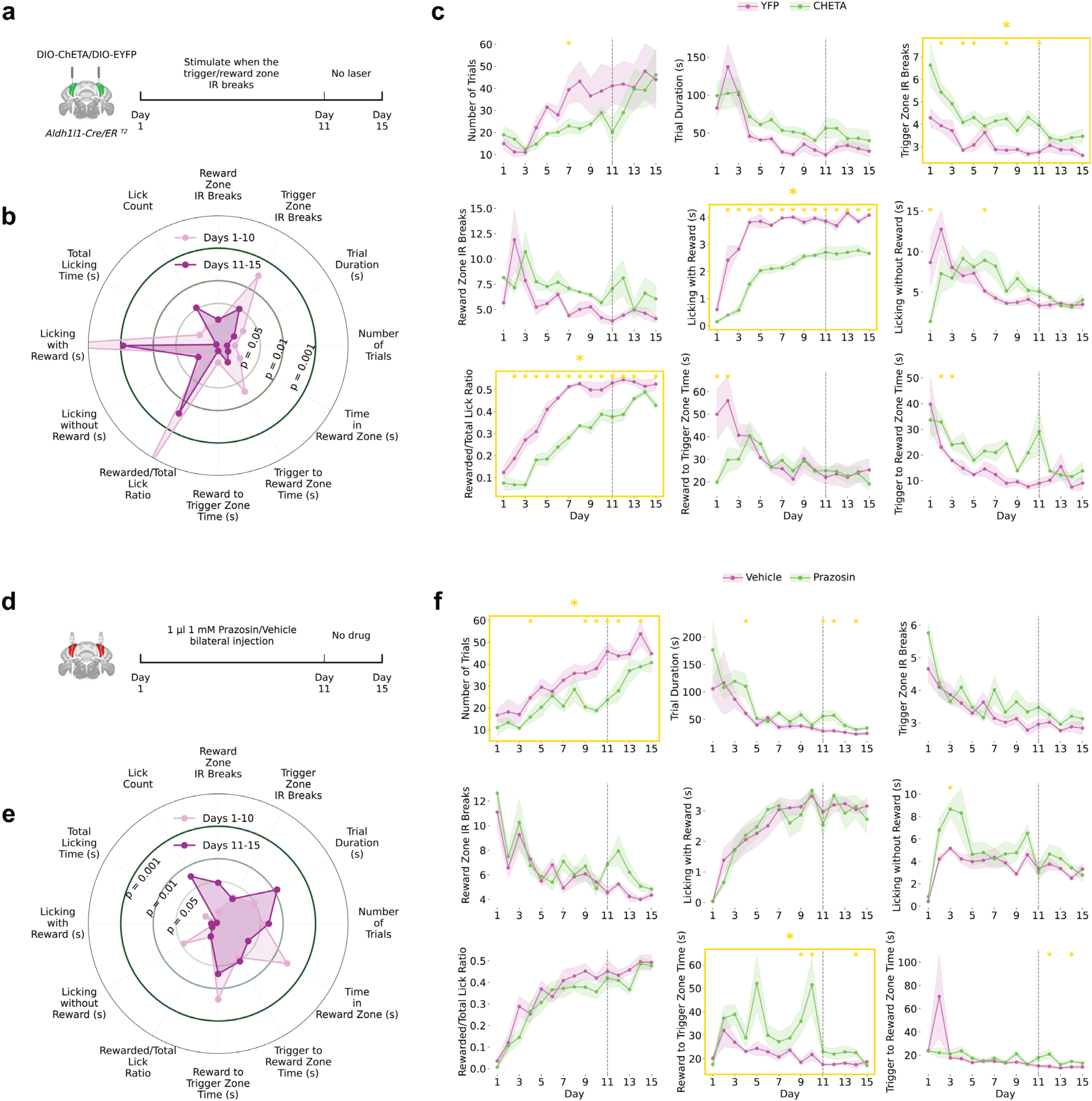
Dissociable Behavioral Roles of Fast and Slow Astrocytic Signals. **(a)** Experimental schematic. ChETA optogenetic stimulation of Bergmann glia was delivered during task events (triggered by IR beam breaks in the trigger/reward zone) in *Aldh1l1-Cre/ER^T^*^2^ mice during Days 1–10, followed by a no-laser period on Days 11–15 (n=6 ChETA, n=5 YFP control). **(b)** Radar plot comparing 10 behavioral metrics between Days 1–10 (light pink) and Days 11–15 (dark purple) in ChETA mice. Concentric rings indicate significance thresholds (p = 0.05, 0.01, 0.001). **(c)** Time-course of behavioral metrics across 15 days in YFP (pink) and ChETA (green) mice. Gold-outlined panels indicate metrics that were significantly different across the entire experimental period; yellow asterisks indicate individual days with significant between-group differences. Shaded regions represent SEM. Dashed line marks the laser-to-no-laser transition. **(d)** Experimental schematic. Bilateral infusion of Prazosin (1 µl, 1 mM) or vehicle was administered daily during Days 1–10, with no drug on Days 11–15 (n=6 Prazosin, n=6 vehicle). Prazosin increased trigger-to-reward and reward-to-trigger transit times and trial duration, and reduced trial number. **(e)** Radar plot comparing behavioral metrics between Days 1–10 and Days 11–15 in vehicle-treated mice. **(f)** Time-course of behavioral metrics across 15 days in Vehicle (pink) and Prazosin (green) mice. Gold-outlined panels indicate metrics significantly different across the entire experimental period; yellow asterisks indicate individual days with significant between-group differences. Shaded regions represent SEM. Dashed line marks the end of drug administration.

ChETA effects depended on task alignment: stimulation delivered every 10 s independently of behavior produced no performance changes (Extended Data Fig. 8), demonstrating that fast BG activity influence requires coupling to task-relevant cues.

For slow signal disruption, the mice received bilateral prazosin or vehicle before daily sessions from Day 1-10, followed by no drug recovery (Day 11-15; Fig. 3d-f). Prazosin impaired distinct behaviors: longer trigger-to-reward time, longer reward-to-trigger time, prolonged trial duration, and reduced trial number—all of which are related to switching between reward-seeking and rewarding states.

BG Ca²⁺ dynamics thus have dissociable roles: fast α2-AR-modulated activities regulate temporally precise cue responses, reinforcement learning, and reward behavior, whereas slow α1-AR-mediated activities support sustained task structure and behavioral transitions.

### Actor-Critic Network Recapitulates Multi-timescale Astrocytic Computation

Our results reveal a division of labor in which astrocytes modulate synaptic transmission at CF and PF inputs to Purkinje cells, differentially regulating instructive versus contextual pathways through distinct temporal modes. Disruption of astrocytic activities impairs learning and performance (Fig. 3), demonstrating that astrocytes actively adjust circuit dynamics to guide behavior.

This modulatory architecture resembles actor-critic frameworks in reinforcement learning^50–53^: actors generate actions, whereas critics evaluate states to modulate learning, improving performance without directly controlling behavior^51^. We hypothesized that this represents a general organizational principle: neurons act as actors generating outputs, whereas astrocytes act as critics evaluating states and modulating computation.

If this captures neuron-astrocyte computation, then artificial networks implementing explicit actor-critic separation should spontaneously develop the multi-timescale, heterogeneous astrocytic dynamics we observe biologically.

We trained a biologically inspired actor-critic network on the spatial sequence task using Proximal Policy Optimization^54,55^ (Fig. 4a). The network comprises 16 neuronal units (actors) generating actions and 16 astrocytic units (critics) estimating state values. Neuron-astrocyte interactions followed tripartite synapse principles^50,56^, where astrocytes bidirectionally modulate synaptic transmission—capturing key properties of biological astrocytic regulation.

**Figure 4:**
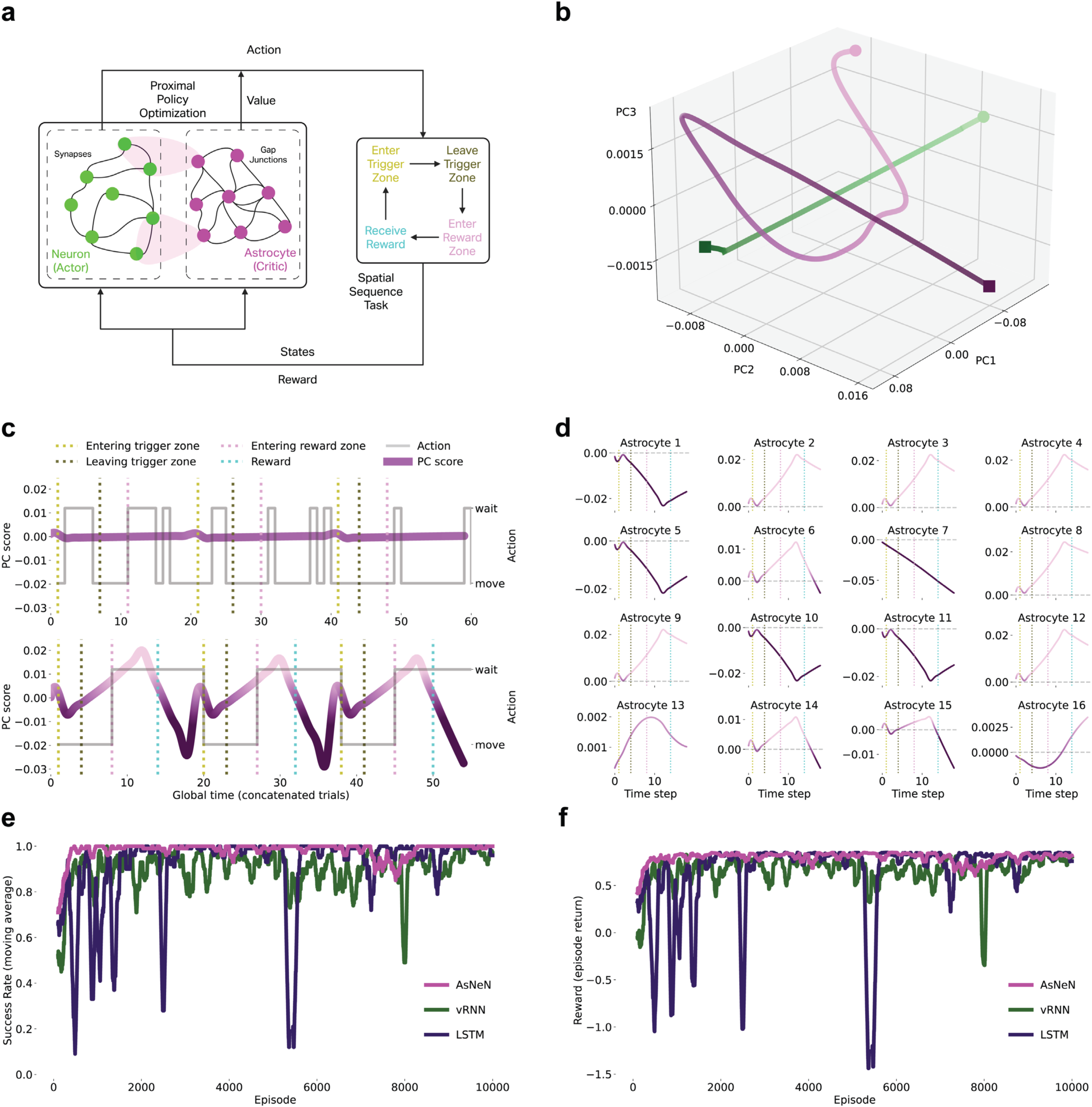
Actor-Critic Network Recapitulates Multi-timescale Astrocytic Computation. **(a)** Biologically inspired actor-critic network with 16 neuronal units (actor) and 16 astrocytic units (critic) implementing tripartite synapse principles. Network trained on the spatial sequence task using PPO. **(b)** PCA of neuronal (green) and astrocytic (magenta) activity showing distinct state-space trajectories. **(c)** Population-averaged astrocytic signals pre- and post-training. Post-training shows fast transients at trigger zone entry and slow ramping signals with polarity reversals resembling biological state-change dynamics. **(d)** Individual astrocyte response profiles across 16 units showing heterogeneous dynamics that recapitulate peak and trough signal diversity. **(e)** Success rate comparison (moving average over 100 episodes) across training for AsNeN (Astrocyte-Neuron Network), vRNN (vanilla Recurrent Neural Networks), and LSTM (Long Short-Term Memory Networks). **(f)** Reward comparison (moving average over 100 episodes) across training for the AsNeN, vRNN, and LSTM networks.

Trained neuronal and astrocytic networks occupied distinct state space regions (Fig. 4b), demonstrating functional differentiation. Critically, multi-timescale astrocytic signals emerged spontaneously through training (Fig. 4c). Pre-training activity was minimal, matching the Day 1 recordings. Post-training revealed fast transients at trigger zone entry (sensory cues) and slow ramping with polarity reversals during state transitions—closely resembling biological state-change dynamics (Fig. 1f).

Individual astrocytes developed heterogeneous response profiles (Fig. 4d), including peak-like and trough-like responses—recapitulating the diversity observed in BG (Fig. 1e, g). The model reproduced the integrative property: individual astrocytes exhibiting peak-like and trough-like signals that, when summed across the population, generated state-change dynamics—providing evidence that state transitions emerge from distributed astrocytic integration.

To assess whether astrocyte-inspired architectures confer computational advantages, we compared our astrocyte-neuron network (AsNeN) with traditional architectures, including variational Recurrent Neural Networks (vRNN) and Long Short-Term Memory networks (LSTM). The AsNeN network outperformed both traditional architectures across multiple performance metrics (Fig. 4e-f). First, the moving-average reward and success rate were consistently higher for AsNeN than for vRNN and LSTM. Second, AsNeN converged faster in terms of both the reward accumulation rate and the success rate. Third, AsNeN exhibited greater stability after convergence, with minimal fluctuations compared with the oscillatory behavior of traditional networks.

Thus, actor‒critic architectures spontaneously develop multi-timescale, heterogeneous astrocytic computations that are observed biologically. Convergence between biological observations and computational theory, combined with superior performance over traditional architectures, suggests that actor-critic computation may be a general organizational strategy for neuron-astrocyte networks during behavior.

## Discussion

### Astrocytes as Multilevel Computational Elements

Our findings establish that astrocytes perform computation by transforming univariate neuromodulatory inputs into multivariate, pathway-specific circuit regulation across distinct timescales. Three features define this computation: temporal decomposition (NE parsed into slow and fast modes through α1-ARs and α2-ARs), spatial specificity (each temporal mode selectively targets distinct synaptic pathways—slow signals modulate PF inputs, fast signals regulate CF inputs), and emergent integration (population-level summation of opposing peak and trough signals generates state-change dynamics). This integrative mechanism receives further support from our actor–critic network simulations, which spontaneously reproduced peak and trough signals that sum to generate state–change dynamics, strengthening the hypothesis that state transitions arise from distributed astrocytic computation^55,56^.

### Complementary Computational Properties

Neurons and astrocytes differ fundamentally in their computational properties. Neurons generate action potentials, providing high-bandwidth, temporally precise, point-to-point signaling^1–3^, whereas astrocytes lack sufficient expression of voltage-gated sodium channels^57,58^ and rely on graded Ca²⁺ elevations^4,5^ that provide lower-bandwidth, temporally extended, spatially integrative signaling. Neurons maintain high membrane resistance for precise signal propagation, whereas astrocytes exhibit low resistance and extensive gap junction coupling^6–8^, generally creating networks that integrate information over large territorial domains and influence tens of thousands of synapses^6–8^. This suggests dual computational architecture: neurons excel at temporally precise, spatially specific computation through electrical signaling; astrocytes excel at temporally extended, spatially integrative computation through chemical signaling^10,59^.

While we focus on norepinephrine, astrocytes express diverse receptors for dopamine, serotonin, and acetylcholine^60–62^, many featuring receptor subtypes with distinct kinetics (e.g., fast nicotinic versus slow muscarinic receptors^63^). If astrocytes simultaneously parse multiple neuromodulators into fast and slow modes, they could generate a combinatorically large control space from few inputs. The fast Ca²⁺ transients we observe may not be unique to Bergmann glia—recent studies identified fast astrocytic signals in the cortex^16–18^, hippocampus^20,21^, and striatum^64^. Our key contribution is that fast signals emerge with learning, show behavioral tuning, and causally regulate behavior. The cerebellar cortex may provide favorable conditions because climbing fibers deliver precisely timed instructive signals^22,23^, but regions with temporally structured inputs may reveal fast signals under appropriate behavioral conditions. Disruption of each timescale results in distinct behavioral deficits—fast signal loss impairs event-triggered responses, whereas slow signal loss impairs state transitions—demonstrating that both modes are necessary and non-redundant.

### Actor-Critic Computation: A Unifying Framework

Our computational modeling revealed that, at multiple timescales, heterogeneous astrocytic dynamics emerge naturally from actor-critic architectures where neurons act as actors generating behavioral outputs and where astrocytes act as critics evaluating states to modulate learning. Implementing neuron-astrocyte interactions following tripartite synapse principles^50,54^ was sufficient to recapitulate biological astrocytic computation, suggesting that bidirectional astrocyte-synapse communication combined with reinforcement learning naturally produces the temporal decomposition and population integration we observe experimentally. The spontaneous emergence of biologically realistic dynamics in artificial networks trained on the same task provides strong evidence that astrocytes perform computation rather than merely reacting to neuronal activity—if astrocytes are passive responders, their dynamics would mirror neuronal patterns rather than developing distinct temporal profiles optimized for state evaluation.

### Implications for Artificial Intelligence

Actor-critic architectures incorporating astrocyte-inspired critic networks can solve complex tasks while spontaneously developing the multi-timescale dynamics observed in biological systems. Our direct comparison demonstrates that the astrocyte-neuron network outperforms traditional architectures, including vRNN and LSTM in learning efficiency, performance, and stability. This suggests design principles for artificial neural networks^65–68^: temporal decomposition layers that automatically parse inputs into fast (event-level) and slow (state-level) components; low-precision, high-integration units with large receptive fields for detecting population patterns^69^; and neuromodulatory decoding modules that transform global signals into pathway-specific regulation. Such architectures could address continual learning challenges by balancing stability (slow-timescale units maintaining the task structure) and plasticity (fast-timescale units enabling rapid adaptation), improve credit assignment through population-level error signals, and enhance meta-learning through three-dimensional control spaces. Recognizing astrocytes as multilevel computational elements may fundamentally reshape the understanding of information processing in biological and artificial systems, potentially leading to more capable artificial systems leveraging complementary computational architectures.

## Methods

### Animals

All procedures were approved by the University of Alabama at Birmingham Institutional Animal Care and Use Committee and followed NIH guidelines. Male and female mice (8-12 weeks old) were group-housed on a 12h light/dark cycle with ad libitum access to food and water except during behavioral testing. The experimental groups included: (i) heterozygous *Aldh1l1*::CreERT2 mice (Jackson Laboratory #029655) crossed with heterozygous *Rosa26*::CAG-GCaMP6f reporter mice for astrocyte-specific calcium ion imaging; (ii) wild-type C57BL/6J mice for GRAB_NE sensor experiments and pharmacological manipulations; (iii) heterozygous *Aldh1l1*::CreERT2 mice for optogenetic manipulations. Tamoxifen (75 mg/kg body weight, i.p.) was administered three times at 2-day intervals between injections to induce Cre recombination in the *Aldh1l1*::CreERT2 lines/crosses. The animals were randomly assigned to experimental groups, and the experimenters were blinded to group assignment during behavioral testing and analysis.

### Viral Constructs and Stereotactic Surgery

For GCaMP6f expression, Aldh1l1-CreERT2; Rosa26-CAG-GCaMP6f mice were used without additional viral injection. For GRAB_NE expression, pAAV_*hSyn*-GRAB_NE1m (Addgene #123308) was injected into cerebellar cortex lobules III and the left and right simplex. For optogenetic experiments, pAAV_*Ef1a*::DIO-ChETA-EYFP or pAAV_ *Ef1a*::DIO-EYFP control (Addgene #26968 and #27056, respectively) was injected into the left and right simplex regions. For plasma membrane Ca^2+^ ATPase (PMCA) overexpression, pZac2.1_*GfaABC1D*::mCherry-hPMCA2w/b (AAV2/5, Addgene#111568) was injected into cerebellar cortex lobules III and left and right simplex.

The mice were anesthetized with isoflurane (1.5-2% in O₂) and placed in a stereotactic frame. Body temperature was maintained at 37°C via a feedback-controlled heating pad. The skull was exposed, and small craniotomies were made above the target coordinates. All coordinates were referenced to the lambda. For cerebellar lobule III, the AP was −2.4 mm from Lambda, the ML was 0 mm, and the DV was 1.0 mm from the brain surface. For the cerebellar simplex, the AP was −2.4 mm from the lambda, the ML was ±2.3 mm, and the DV was 1.0 mm from the brain surface. The virus (200 nl per site) was injected at 50 nl/min using a micropump, and the needle remained in place for 10 min post-injection before slow retraction. For fiber photometry, fiber optic cannulas (200 μm core diameter, 0.37 NA, RWD Life Science) were implanted above the injection sites and secured with dental cement. For pharmacological experiments, guide cannulas (RWD Life Science) were implanted bilaterally at target sites. For combined photometry and drug delivery, multiple fluid injections cannulas (200 μm, 0.37 NA, Doric Lenses) were used. The animals were allowed to recover for 3‒4 weeks before behavioral training to allow optimal expression.

### Behavioral Task

Mice were water restricted to 1 ml per day starting 2 days before the behavioral experiments and trained on a custom spatial sequence task in a rectangular arena (60 cm × 8 cm) with distinct trigger and reward zones (10 cm × 10 cm each) at opposite ends. Infrared (IR) beam-break sensors detected zone entries/exits. The task required: (1) entering the trigger zone, which activated an auditory tone (8 kHz, 0.2 s duration); (2) exiting the trigger zone and traversing to the reward zone; (3) entering the reward zone and waiting 2 s, after which a liquid sucrose reward (5% w/v, 4 μl) was delivered via a solenoid-controlled lick port; (4) consuming reward during a 4 s window; and (5) returning to the trigger zone to initiate the next trial. Lick events were detected by the breaking of the IR sensor at the port. Training consisted of 30 min daily sessions for 15 consecutive days. Day 1 served as a no-reward baseline to assess exploratory behavior. The following behavioral metrics were used: number of successful trials, trigger-to-reward zone time, reward-to-trigger zone time, trial duration, licking bouts with/without a reward, and the rewarded licks/total licks ratio.

### Fiber Photometry

Fiber photometry was performed using an RWD R821/FR-21 Tricolor Multichannel Fiber Photometry System. Both blue LED light (470 nm) and an isosbestic wavelength control (410 nm) were modulated at 15 Hz during recording and delivered through the implanted fiber. The signals were digitized at 1 kHz and synchronized with behavioral events via TTL pulses from IR sensors.

The data were analyzed in MATLAB and Python. For the GCaMP6f signals, raw fluorescence (F) was used to calculate ΔF/F = (F_470_ − F_410_)/F_410_ after photobleaching was corrected via exponential fitting. For GRAB_NE signals, the 0-200 s period of each session was used as a baseline for normalization, and then a −2 to +2 s window around each event was used for event-specific baseline correction. The signals were then normalized to z-scores for cross-session comparisons. The training data were defined as the day the animal completed the most trials after day 7.

Slow signal classification was performed based on activity patterns during licking onset. Recording sites were classified as trough, state-change, or peak signals based on the presence of troughs or peaks within a −1 to +1 s window around licking onset. Sites showing clear troughs were classified as trough signals, sites showing clear peaks were classified as peak signals, and sites showing neither prominent peaks nor troughs were classified as state-change signals.

### Pharmacology

For receptor antagonism during photometry recordings, drugs were infused through the multiple fluid injections cannulas (Doric Lenses) used for combined optical recording and drug delivery after 10 days of training. Yohimbine hydrochloride (α2-AR antagonist, Tocris), prazosin hydrochloride (α1-AR antagonist, Tocris), propranolol hydrochloride (β-AR antagonist, Tocris), or vehicle (artificial cerebrospinal fluid, aCSF) were prepared at 1 mM concentration and infused at 1 μl per recording site at 250 nl/min. After infusion completion, the injector remained in place for 5 min to prevent backflow. Behavioral sessions and recordings began 10 min post-infusion. Each animal received all the treatments in counterbalanced order, with ≥3 days washout between sessions.

For behavioral manipulation, prazosin (1 mM in aCSF, 1 μl per side) or vehicle was infused bilaterally 10 min before daily training sessions (Days 1-10) via implanted guide cannulas using the same infusion protocol (250 nl/min, 5 min hold time). Infusions ceased on Days 11-15 to assess recovery.

### Optogenetic Manipulation

For fast signal disruption, *Aldh1l1*::CreERT2 mice expressing ChETA-EYFP or control EYFP in Bergmann glia received bilateral fiber optic implants above the cerebellar simplex. Laser stimulation (473 nm, 10 mW, 50 ms pulses) was triggered automatically by IR beam breaks when the animals entered or exited the trigger zone or reward zone during Days 1-10 of training. To prevent repeated stimulation during prolonged zone occupancy, a 5 s refractory period was implemented following each stimulation event. Stimulation ceased on Days 11-15 to assess recovery. Control experiments used the same stimulation protocol but delivered independently of behavior (every 10 s) or used EYFP-expressing control animals with behavior-triggered stimulation.

### Ex Vivo Electrophysiology

Acute cerebellar slices (300 μm thick) were prepared from wild-type mice (3-8 weeks old) or mice expressing PMCA (4 weeks post-AAV injection). The mice were deeply anesthetized with a ketamine and xylazine mixture and transcardially perfused with ice-cold cutting solution containing (in mM): 87 NaCl, 2.5 KCl, 0.5 CaCl₂, 7 MgCl₂, 1.25 NaH₂PO₄, 25 NaHCO₃, 25 glucose, and 75 sucrose, bubbled with 95% O₂/5% CO₂. The brain was rapidly removed and cut transversely using a vibratome (VT1200S, Leica Microsystems). Slices were transferred to oxygenated aCSF at 32°C for 30 min and then allowed to recover for 1 h at room temperature (25 °C) before recordings in aCSF containing (in mM): 119 NaCl, 2.5 KCl, 2.5 CaCl₂, 1.3 MgCl₂, 1.3 NaH₂PO₄, 26 NaHCO₃, and 20 glucose.

Whole-cell voltage-clamp recordings were obtained from visually identified Purkinje cells using an upright microscope (Axio Examiner. D1, Zeiss) with IR-differential interference contrast (DIC) optics. Individual slices were transferred to a submerged chamber and continuously perfused with normal oxygenated aCSF at room temperature. Patch pipettes (3-4 MΩ) were filled with internal solution containing (in mM): 120 Cs-gluconate, 17.5 CsCl, 10 Na-HEPES, 4 Mg-ATP, 0.4 Na-GTP, 10 Na₂-creatine phosphate, and 0.2 Na-EGTA (290-300 mOsm, pH 7.3). Cells were held at −60 mV. Cells with serial resistances above 25 MΩ were discarded, and cells were also excluded if any whole-cell parameter (i.e., Cm, Ri, Rs) changed by ≥20% during recordings.

For climbing fiber and parallel fiber stimulation, aCSF-filled pipettes connected to an isolated stimulator (ISO-Flex, AMPI) were used to deliver electrical stimulation every 20 s. For climbing fiber stimulation, the stimulating electrode was placed in the granule cell layer to elicit all-or-none climbing fiber excitatory post-synaptic potentials (EPSCs). For parallel fiber stimulation, the electrode was placed in the molecular layer to evoke graded EPSCs. The stimulus intensity was adjusted to ensure that EPSCs were clamped. Synaptic responses were recorded at baseline (0-300 s), during drug application (300-1100 s), and during washout (1100-1880 s). Prazosin (10 μM) or yohimbine (15 μM) was bath applied during the drug application period.

Data were acquired using TI Workbench software with a MultiClamp 700B amplifier (Molecular Devices), filtered at 2 kHz, and digitized at 10 kHz with ITC-18 A/D-D/A interfaces (Instrutech). Analysis was performed using TI Workbench and Python.

### Immunohistochemistry

Mice were deeply anesthetized and transcardially perfused with 4% (w/v) paraformaldehyde (PFA) in phosphate-buffered saline (PBS). Brains were removed and post-fixed overnight in 4% PFA at 4°C. Coronal brain sections were cut at the thickness of 60 μm using a vibratome. Sections were permeabilized with 0.25% (v/v) Triton X-100 for 2 h at room temperature and blocked with 10% (v/v) normal goat serum for 1 h. Sections were incubated at 4°C overnight with blocking solution containing the following primary antibodies: rabbit anti-α1-AR (1:500, Invitrogen), rabbit anti-α2-AR (1:500, Proteintech), mouse anti-S100β (1:500, Invitrogen), and guinea pig anti-VGLUT1 (1:500, MilliporeSigma). After primary antibody incubation, the sections were rinsed with PBS three times for 10 min each and incubated for 2 h at room temperature with Alexa Fluor™-conjugated secondary antibodies (1:500, Jackson ImmunoResearch Laboratories). The sections were coverslipped with Vectashield anti-fade mounting medium (Vector Laboratories). Confocal images were acquired on a Zeiss LSM 880 microscope using 20 x (for overview) and 40x oil-immersion objectives (to obtain cellular detail). Colocalization analysis was performed using ImageJ with the JACoP plugin.

### Bio-Inspired Neuron-Astrocyte Network Simulation

To simulate and investigate how astrocyte activity supports and modulates behavior, we designed an artificial task in which an agent equipped with a biologically inspired neuron–astrocyte network is trained with reinforcement learning (RL) to solve the task in a manner analogous to the experiment.

#### Task environment: spatial sequence reward task

We built an artificial spatial sequence reward task that captures the key structure of the mouse experiment. In the task, the agent needs to move from a trigger zone to a reward zone and obtain a reward after a delay. Each trial is implemented as a discrete-time Markov decision process with four states: a trigger zone, a transit state, a reward zone, and the terminal success or failure. At each time step, the agent can select one of two actions, move forward or wait/stay.

At the beginning of a trial, the agent typically starts in the trigger zone, where a brief cue is initiated, representing the sensory cue in the experiment. If the agent chooses the move action, the state transitions into the transit phase, which lasts a fixed number of time steps. The agent needs to keep moving in this phase and then can enter the reward zone. In the reward zone, the agent must emit the wait action for enough consecutive time steps to receive the reward. This sustained waiting period mimics licking at the reward spout after a fixed delay. When the accumulated hold time exceeds the reward delay, the agent receives a unit reward (+1), followed by a brief post-reward period, after which the episode terminates in a successful state. If the agent fails to obtain a reward before a hard maximum trial length, the episode terminates in failure. During the task, each time step has a small step cost (−0.01), so longer or inefficient trajectories are mildly penalized, encouraging the agent to move efficiently and to wait only as long as necessary.

At each time step, the agent receives a three-dimensional observation: the first component is a constant bias, i.e., +1; the second is the cue channel, which is transiently active after entering the trigger zone, i.e., +1 or 0; and the last component indicates whether the reward has been delivered, i.e., +1 or 0. This minimal environment preserves the temporal structure of the experiment, namely, cue-gated departure from the trigger zone, delayed reward contingent on sustained waiting, and time-limited trials. Moreover, it remains simple enough and facilitates a systematic analysis of learned neuron–astrocyte dynamics.

#### Bio-inspired neuron-astrocyte network

The agent was implemented with a biologically inspired recurrent neuron-astrocyte network with standard linear input and output layers. The recurrent network is an extension of the previous neuron-astrocyte network model^54^, which was built based on the tripartite synapse structure of neuron‒astrocyte interactions. The network consists of neuronal activities x_t_ ∈ ℝ^{Nₓ}, astrocytic activities z_t_ ∈ ℝ^{N_z_}, and a flattened synaptic weight vector w_t_, which is reshaped into a recurrent weight matrix W_t_. In the simulations reported here, we set Nₓ = N_z_= 16, and the synaptic weight w_t_ is sparsely initialized.

Let o_t_ denote the 3-dimensional observation from the task environment at time t. The updates for neurons, synapses, and astrocytes are given by the following system:

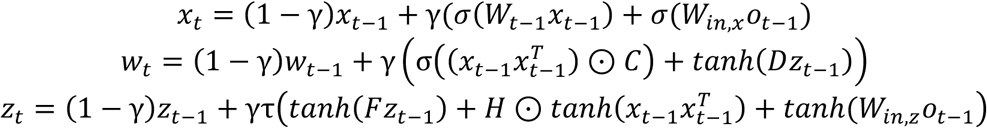

Here, σ is the sigmoid activation, and tanh is the hyperbolic tangent activation function. γ is the fixed discretization step, and τ < 1 denotes the slow astrocyte dynamics relative to neurons. The matrix C parameterizes element-wise gains on the outer product x_t_ x_t_^T^, D maps astrocyte activity to synaptic modulation, H maps synaptic activity back to astrocytes, and F captures astrocyte–astrocyte coupling. The matrices W_in,x_ and W_in,z_ inject task observations into neurons and astrocytes, respectively. This architecture implements the biologically tripartite feedback loop in which neuronal activity drives synaptic and astrocytic updates, and astrocytes feedback to synapses and neurons to provide slow, context-dependent modulation (see more details of the network model in the early publication^54^).

#### Reinforcement learning agent and training procedure

We implemented this neuron-astrocyte-network-embedded agent in the reinforcement learning framework, i.e., the proximal policy optimization (PPO), to solve the defined spatial sequence reward task. We define separate readouts to generate the policy and value functions. For the policy, we interpret the neuronal layer as the action-selecting module but allow astrocytes to modulate it. Specifically, the action probabilities over the two discrete actions (move, wait) are given by a SoftMax readout π(a_t_ | o_t_) = SoftMax(W_π x_t_). The value function is output directly from astrocyte activity via a linear map V(o_t_) = W_v z_t_, reflecting the role of astrocytes as a slower critic-like integrator of reward history and task context.

All the parameters in the neuron-astrocyte network, policy and value readouts are trained with the Adam optimizer in Python using multiple PPO epochs per update and minibatches. During the training, we track the obtained reward and the binary success flag for each episode. To assess the convergence and performance, we compute the average reward and success rates, respectively, over a moving window of multiple episodes. All simulations are implemented in Python using PyTorch with fixed random seeds and deterministic settings to enhance reproducibility. The settings and hyperparameters used in the simulation are summarized in Table 1.

**Table 1.** Simulation settings.

| Category | Parameter | Value |
| --- | --- | --- |
| Network | Neuron units ($N_x$) | 16 |
| | Astrocyte units ($N_z$) | 16 |
| | Discretization step ($\gamma$) | 0.01 |
| PPO / RL | Policy learning rate | 0.001 |
|  | PPO clip | 0.2 |
|  | Entropy coefficient | 0.05 |
|  | Rollout horizon (T) | 64 |
|  | PPO epochs per update | 16 |
|  | Minibatch size | 64 |
|  | Training episodes | 10,000 |
| Environment | Reward delay | 2-8 |
|  | Maximum trial length | 20 steps |

#### Analysis of trained network dynamics

After training, we analyze both the behavioral performance of the agent and the internal dynamics of the neuron–astrocyte network, focusing on astrocyte activity and its interaction with neuronal activity. We generate rollouts using the trained policy in a fixed environment configuration. For each time step, we record the full internal state [x_t_, w_t_, z_t_], the environment state label, the chosen action, and key task events (i.e., entering the trigger zone, leaving the trigger zone, entering the reward zone, and receiving the reward).

We separated the recorded states into neuronal activity x_t_ and astrocyte activity z_t_ and examined their dynamics in single and multiple successful trials. To obtain low-dimensional summaries, we perform standard principal component analysis (PCA) on x_t_ and z_t_. For single trials, PCA is computed on that trial’s activity; for multi-trial analyses, we concatenated activity across several successful trials, computed a shared PCA basis, and then projected each trial into this basis. We plotted the first principal component as a function of time, aligned to behavioral events and overlaid with the action sequence. To visualize the full astrocyte population, we also plotted the raw states of individual astrocytes across time and units. These visualizations reveal both fast, event-locked components and slower ramps or plateaus aligned to reward-related epochs, which aligns with the experimental findings.

As a control, we also performed the same analyses on untrained (randomly initialized) networks underlying the same environment. We compared trained and untrained dynamics, which demonstrate how neuronal and astrocytic activities are reshaped in the reinforcement learning process to solve the task.

#### Learning performance comparison with standard network architectures

We further evaluated the bio-inspired neuron-astrocyte network by comparing it with two standard recurrent architectures, a vanilla RNN and an LSTM, applied to the same artificial task. Both baseline networks use a hidden dimension of 128, with a parameter count comparable to that of the neuron-astrocyte network. All the networks were trained using the same PPO framework with the same hyperparameter settings. Then, we compared the learning performance by monitoring two metrics: the episodic reward and the success rate, each reported as a moving average over a 100-episode window. The results showed that the neuron-astrocyte network outperforms these two networks by having relatively higher reward and success rates throughout training. Moreover, its learning curves converge faster and exhibit much less variance, which indicates better learning stability than the standard recurrent baselines do.

### Statistical Analysis

All the statistical analyses were performed in MATLAB, Python, and GraphPad Prism. The data are presented as the means ± SEMs unless otherwise noted. Sample sizes were determined based on pilot experiments and previously published studies; sample sizes are consistent with standards in the field The normality of the data was assessed using Shapiro-Wilk test. For normally distributed data, parametric tests were used: paired or unpaired t tests for two-group comparisons and one-way or two-way ANOVA with Tukey’s or Sidak’s post hoc tests for multiple comparisons. For non-normally distributed data or small sample sizes, non-parametric tests were used: Mann-Whitney U test, Wilcoxon signed-rank test, or Kruskal-Wallis test with Dunn’s post hoc test. Repeated-measures ANOVA was used for longitudinal behavioral data across training days. Correlations were assessed using Spearman’s rank correlation coefficient for non-parametric data or when monotonic relationships were examined. Statistical significance was set at α = 0.05.

## Data and Code Availability

All data supporting the findings of this study and custom analysis code are available from the corresponding author upon reasonable request. The computational models will be deposited in a public repository (GitHub) upon publication.

## Author Contributions

CL: conceptualization, investigation, formal analysis, visualization, project administration, writing – original draft, and writing – review & editing. LG: methodology, formal analysis, software, validation, visualization, and writing – review & editing. CS: software, data curation, formal analysis, and writing – review & editing. SC: methodology and writing – review & editing. LP-M: conceptualization, funding acquisition, resources, supervision, and writing – review & editing. WL: conceptualization, funding acquisition, resources, supervision, project administration, and writing – review & editing.

## Acknowledgements

This work was supported by the National Institutes of Health grants R01-NS121542, R21-NS108508, and R21-NS120315. We thank Dr. Vladimir Parpura for helpful comments. We are grateful to Yijian Zhang, Likhitha Polepalli, and Jenny Shen for maintaining the mouse colony. We also thank lab members Julia Lopes Goncalez, César Acevedo-Triana, Xin Xu, Destynie Medeiros, Suraj Cherian, and Akash Saxena for their assistance and support.

---

**Extended Data Figure 1:**
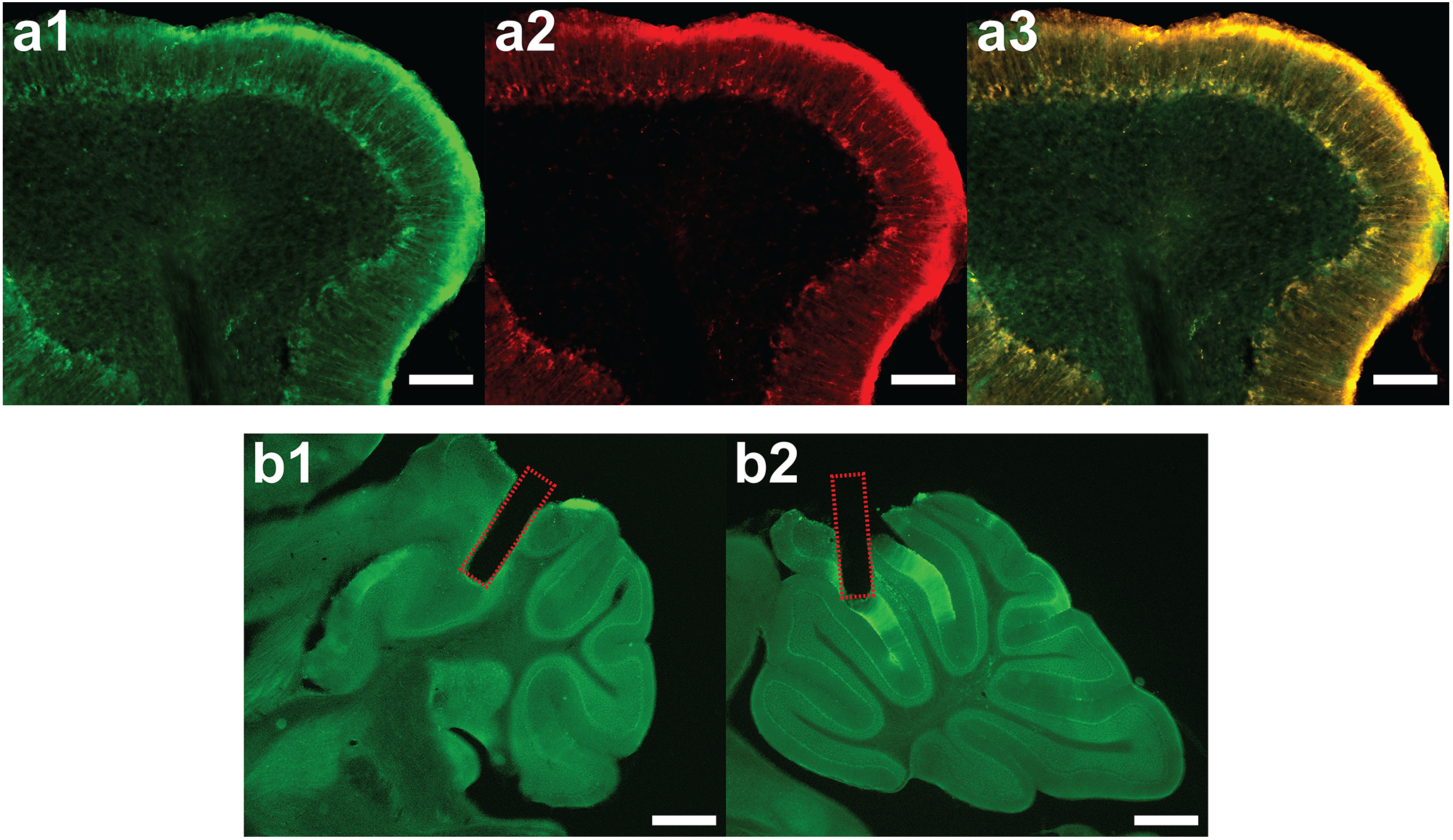
Validation of Bergmann Glia Expression and Fiber Placement. **(a1)** GCaMP6f immunostaining (green) showing expression in Bergmann glia. Scale bar: 100 μm. **(a2)** S100β astrocyte marker immunostaining (red). Scale bar: 100 μm. **(a3)** Colocalization of GCaMP6f and S100β, confirming astrocyte-specific expression. Scale bar: 100 μm. **(b1)** Fiber photometry cannula placement in left and right simplex. Scale bar: 1000 μm. **(b2)** Fiber photometry cannula placement in midline lobule III. Scale bar: 1000 μm.

**Extended Data Figure 2:**
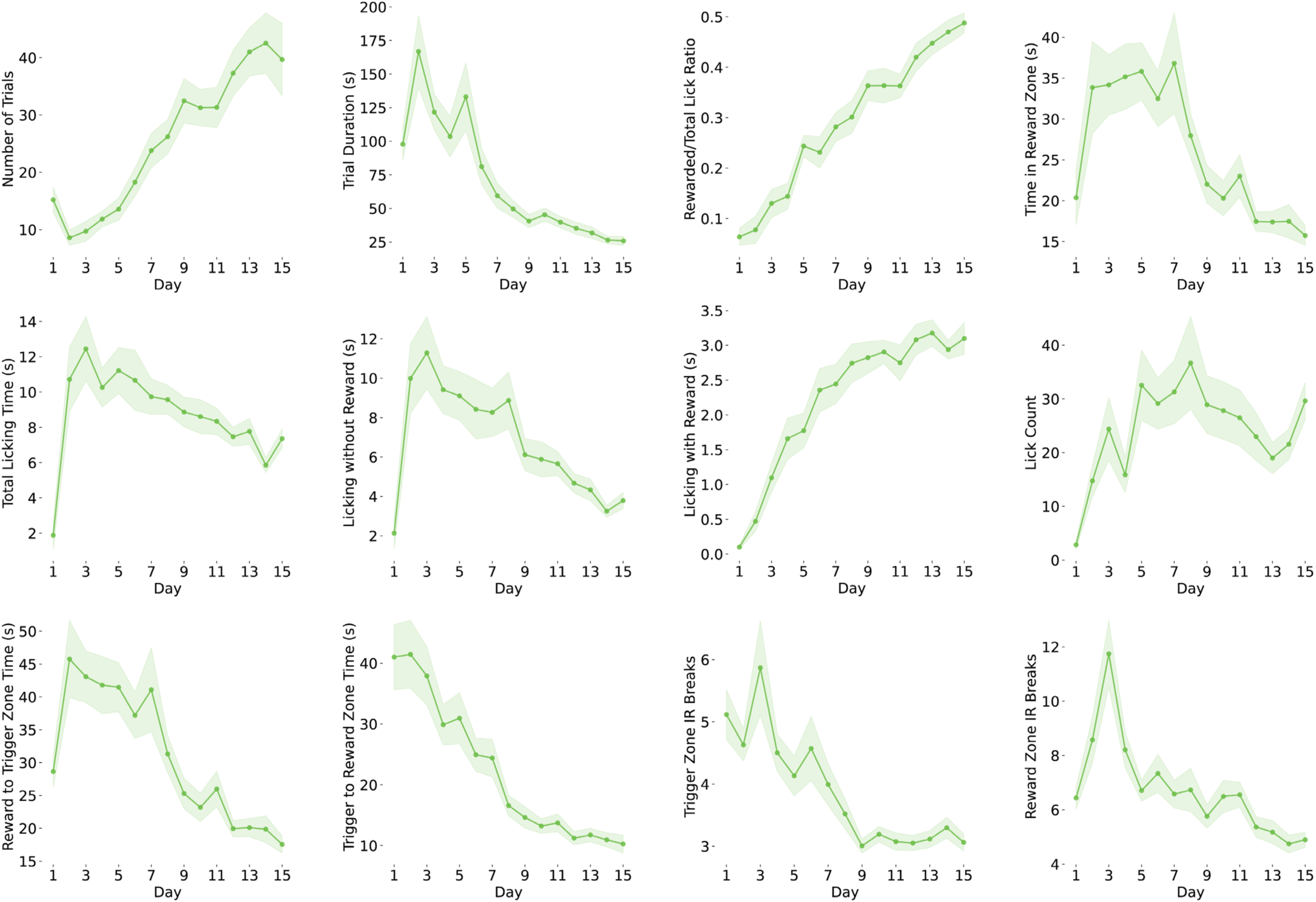
Behavioral Learning Across Training. Training-dependent improvements (Days 1-15) in trial number, trial duration, rewarded/total lick ratio, reward zone time, total licking time, licking with/without reward, licking counts, trigger zone to reward zone time, reward zone to trigger zone time, and trigger zone/reward zone IR breaks.

**Extended Data Figure 3:**
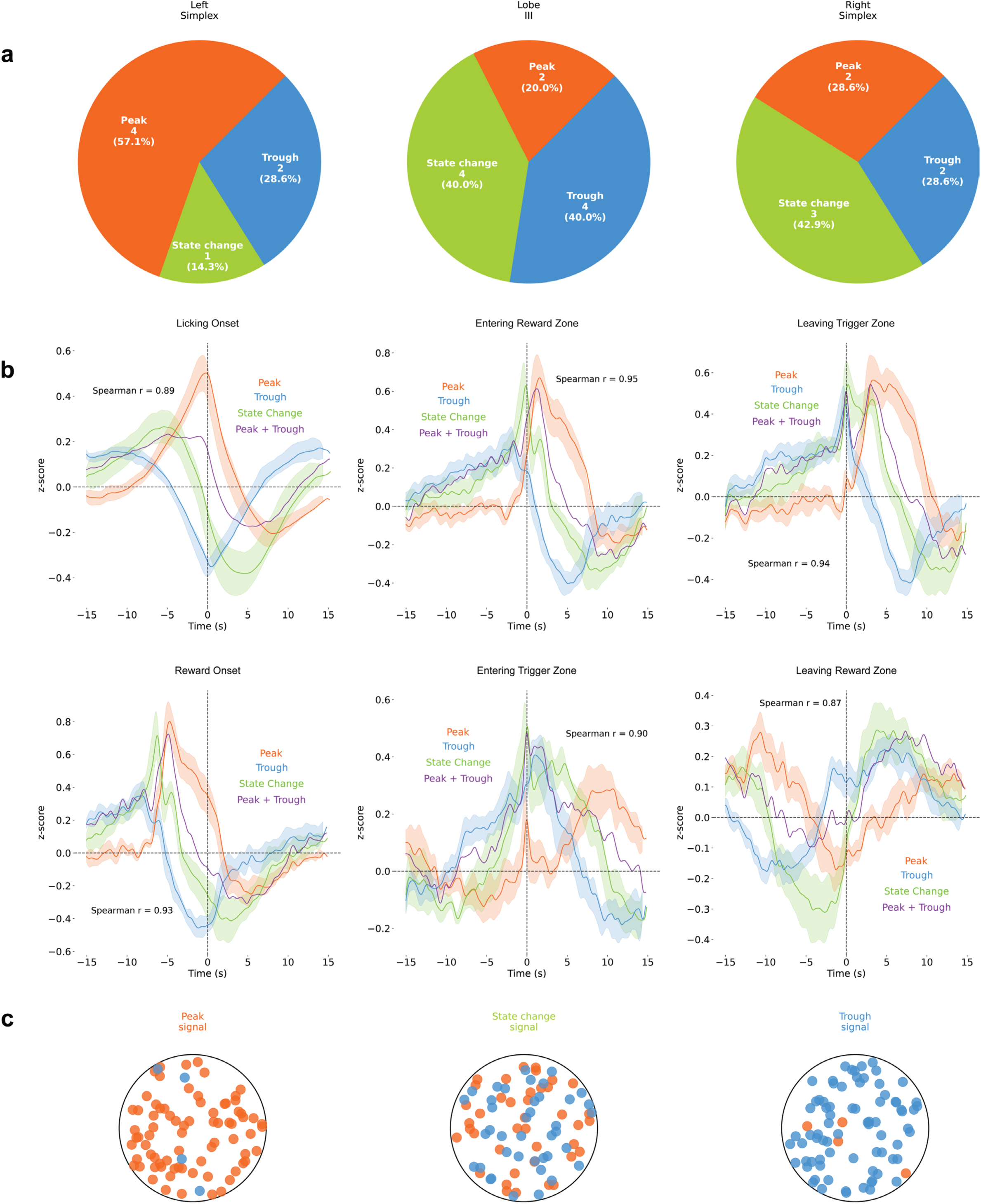
Spatial Distribution of Slow Signal Types Across Cerebellar Regions. **(a)** Signal type distribution across the left simplex, lobule III, and right simplex shows no clear anatomical bias. **(b)** The sum of the peak and trough signals (z-scored ΔF/F) approximates the state-change dynamics (Spearman r = 0.87-0.95), suggesting population integration. **(c)** Schematic of heterogeneous signal types within photometry field of view.

**Extended Data Figure 4:**
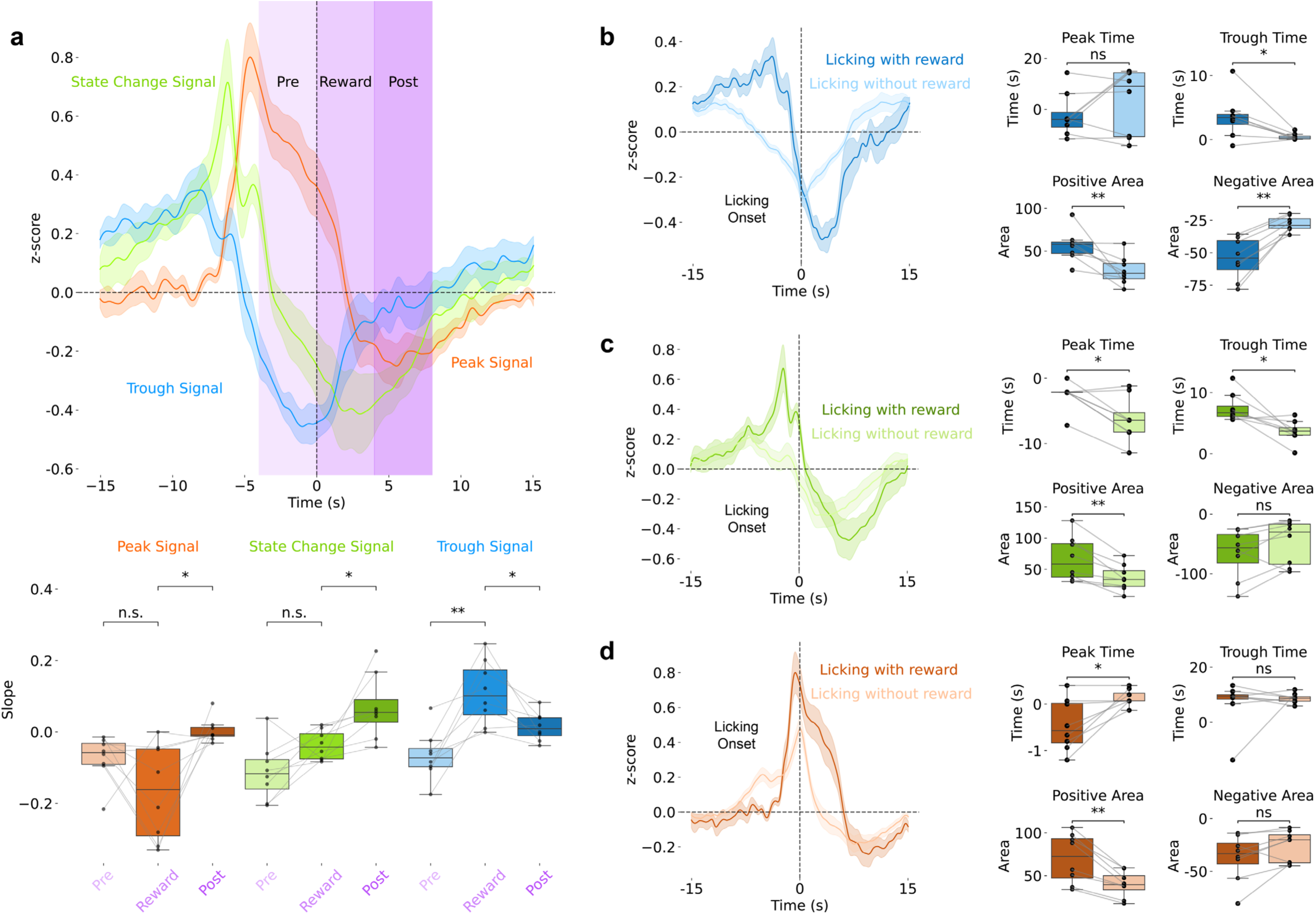
Slow Ca²⁺ Signal Dynamics Correlated with Reward. **(a)** Slow signal types (z-scored ΔF/F) aligned to reward onset with slope quantification across pre-reward (−4 to 0 s), reward (0 to 4 s), and post-reward (4 to 8 s) epochs. **(b-d)** Rewarded versus unrewarded licking comparison (z-scored ΔF/F) showing differential dynamics in the **b)** trough, **(c)** state-change, and **(d)** peak signals.

**Extended Data Figure 5:**
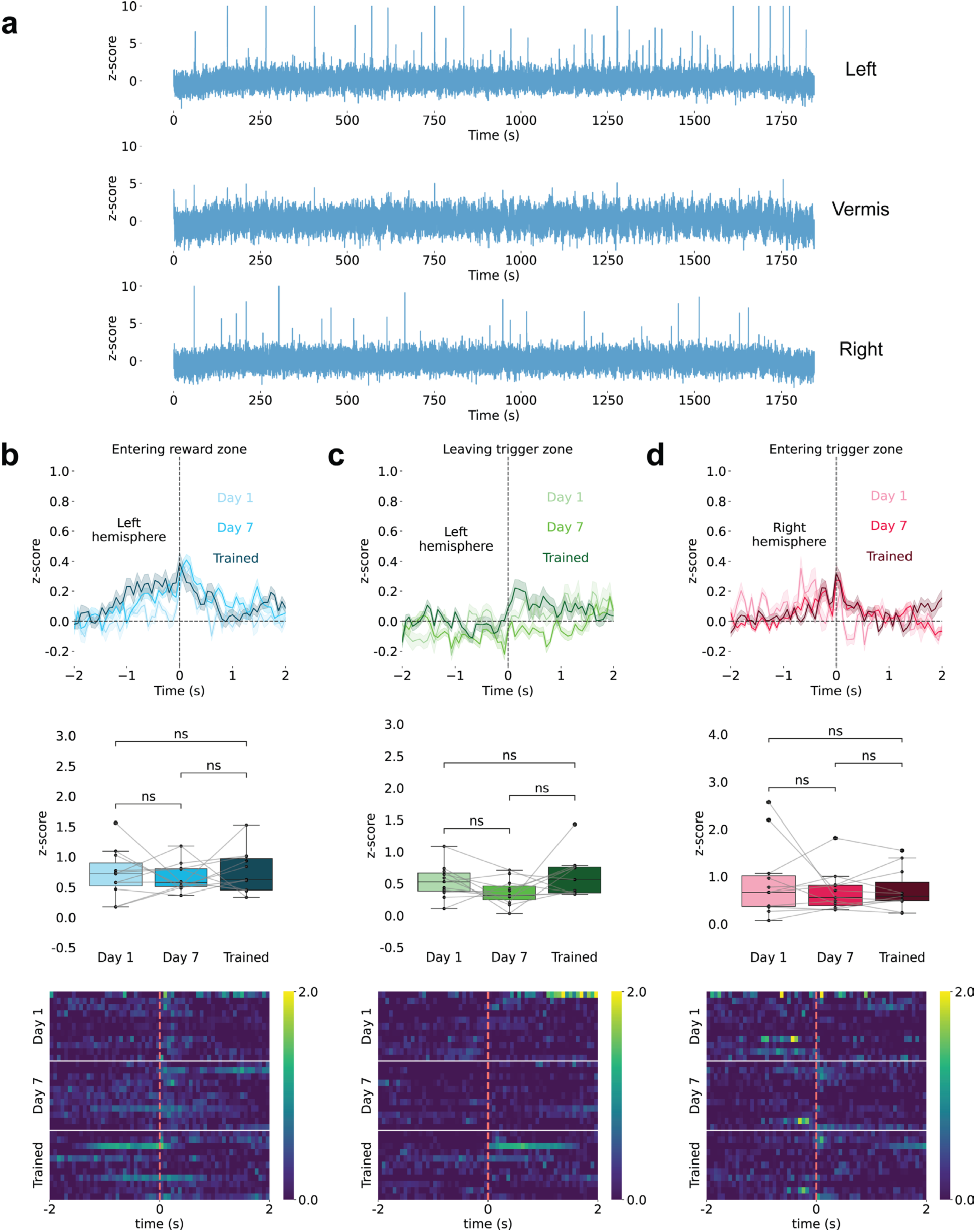
Raw Traces Showing Fast Transients Across Recording Regions. **(a)** Fast transients (z-scored ΔF/F) in the lateral simplex but not midline lobule III. **(b-d)** Lobule III shows no significant fast responses (z-scored ΔF/F) to **(b)** entering reward zone, **(c)** leaving trigger zone, or **(d)** entering trigger zone across training (n=11 sites, 11 animals).

**Extended Data Figure 6:**
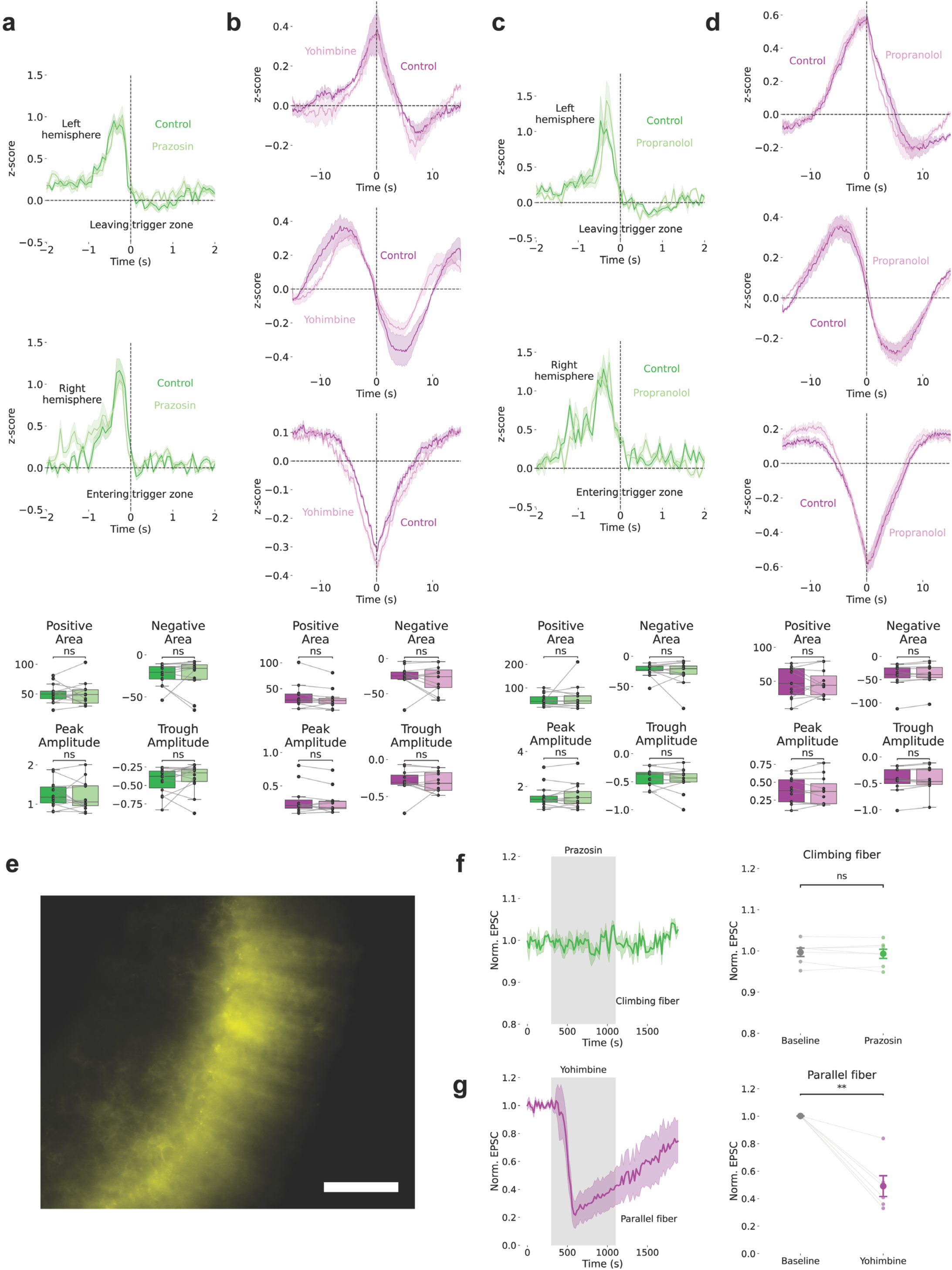
Pharmacological Specificity of Adrenergic Receptor Effects. **(a)** Prazosin (an α1-AR antagonist) does not affect fast transients (z-scored ΔF/F). **(b)** Yohimbine (an α2-AR antagonist) does not affect slow signals (z-scored ΔF/F). **(c-d)** Propranolol (β-AR antagonist) does not affect slow or fast signals (z-scored ΔF/F). **(e)** PMCA expression in Bergmann glia. Scale bar: 50 μm. **(f)** Prazosin does not affect CF-evoked EPSCs. **(g)** Yohimbine reduces PF-evoked EPSCs.

**Extended Data Figure 7:**
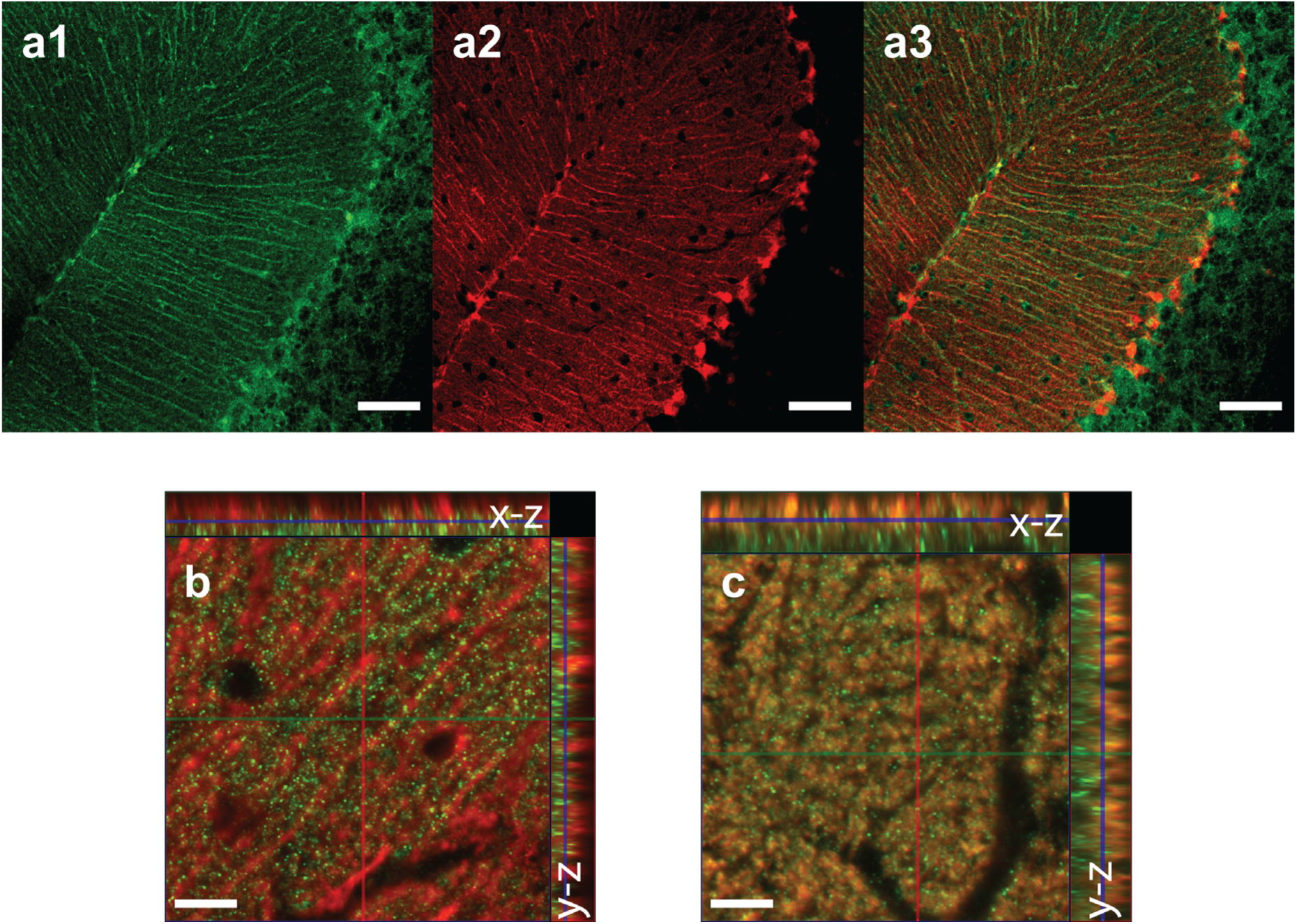
Anatomical Localization of Adrenergic Receptors. **(a1)** α2-AR immunostaining showing expression in cerebellar cortex Bergmann glia molecular layer. Scale bar: 50 μm. **(a2)** S100β astrocyte marker immunostaining. Scale bar: 50 μm. **(a3)** Colocalization of α2-AR and S100β. Scale bar: 50 μm. **(b)** Colocalization of α1-AR and S100β astrocyte marker. Scale bar: 20 μm. **(c)** Colocalization of α1-AR and VGLUT1-positive parallel fiber terminals. Scale bar: 20 μm.

**Extended Data Figure 8:**
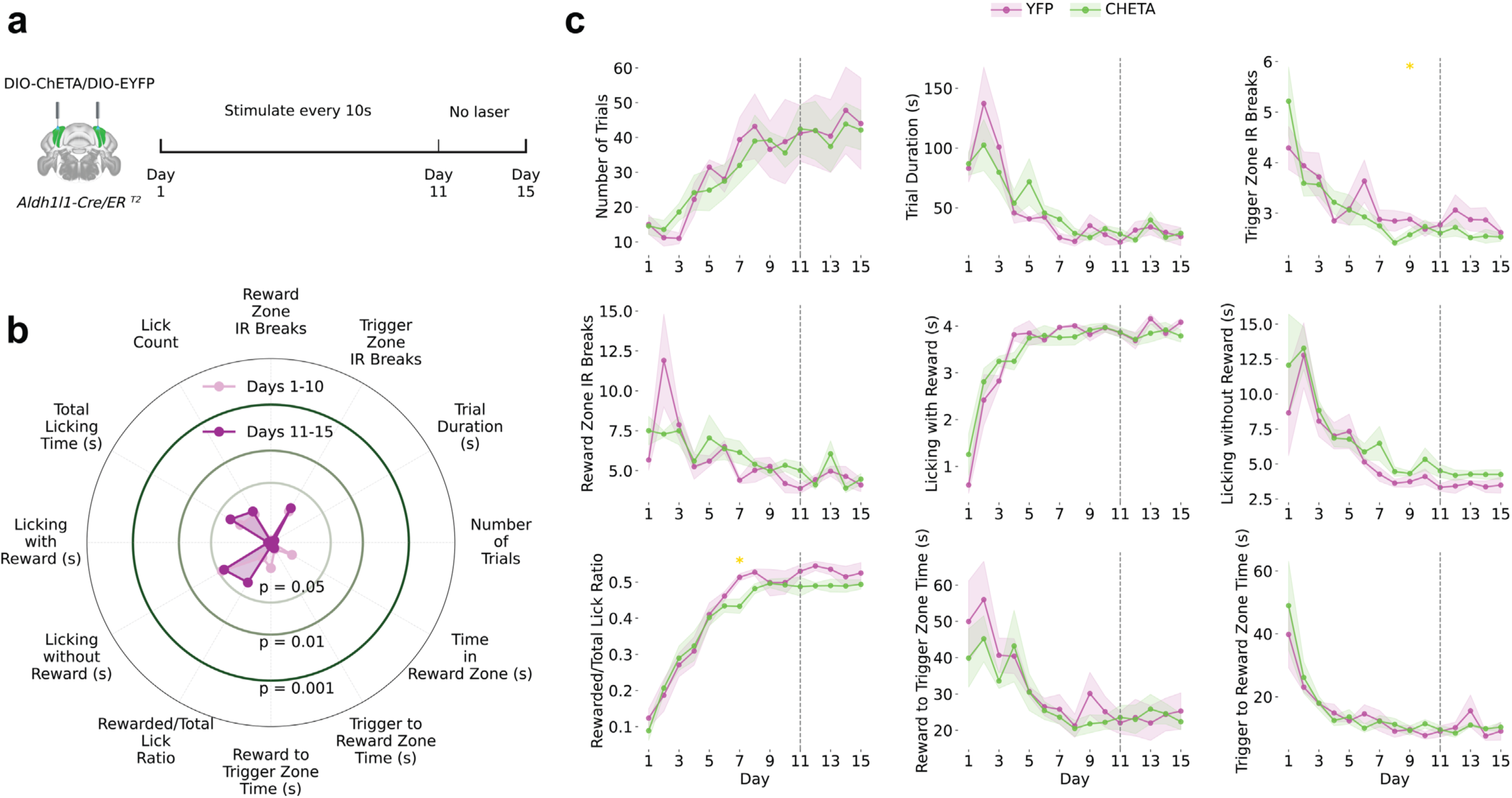
Task-Independent Effects of Optogenetic Fast Signal Disruption. **(a)** Non-contingent control experiment. ChETA optogenetic stimulation of Bergmann glia was delivered every 10 seconds independent of behavior in *Aldh1l1-Cre/ER^T^*^2^ mice during Days 1–10, followed by a no-laser period on Days 11–15 (n=7 ChETA, n=5 YFP control). **(b)** Radar plot comparing 10 behavioral metrics between Days 1–10 (light pink) and Days 11–15 (dark purple) in ChETA mice. Concentric rings indicate significance thresholds (p = 0.05, 0.01, 0.001). **(c)** Time-course of behavioral metrics across 15 days in YFP (pink) and ChETA (green) mice. Yellow asterisks indicate individual days with significant between-group differences. Shaded regions represent SEM. Dashed line marks the laser-to-no-laser transition. Task-independent stimulation produced no significant behavioral changes compared to YFP controls, demonstrating that disruption of Bergmann glia signaling requires temporal coupling to task-relevant cues and is not attributable to nonspecific optogenetic activation.

## References

1. Hodgkin AL, Huxley AF. A quantitative description of membrane current and its application to conduction and excitation in nerve. J Physiol. 1952;117(4):500–544.

2. McCulloch WS, Pitts W. A logical calculus of the ideas immanent in nervous activity. Bull Math Biophys. 1943;5:115–133.

3. Carandini M, Heeger DJ. Normalization as a canonical neural computation. Nat Rev Neurosci. 2012;13(1):51–62.

4. Bazargani N, Attwell D. Astrocyte calcium signaling: the third wave. Nat Neurosci. 2016;19(2):182–189.

5. Volterra A, Liaudet N, Savtchouk I. Astrocyte Ca²⁺ signalling: an unexpected complexity. Nat Rev Neurosci. 2014;15(5):327–335.

6. Bushong EA, Martone ME, Jones YZ, Ellisman MH. Protoplasmic astrocytes in CA1 stratum radiatum occupy separate anatomical domains. J Neurosci. 2002;22(1):183–192.

7. Halassa MM, Fellin T, Haydon PG. The tripartite synapse: roles for gliotransmission in health and disease. Trends Mol Med. 2007;13(2):54–63.

8. Araque A, Parpura V, Sanzgiri RP, Haydon PG. Tripartite synapses: glia, the unacknowledged partner. Trends Neurosci. 1999;22(5):208–215.

9. Paukert M, Agarwal A, Cha J, et al. Norepinephrine controls astroglial responsiveness to local circuit activity. Neuron. 2014;82(6):1263–1270.

10. Savtchouk I, Volterra A. Gliotransmission: Beyond black-and-white. J Neurosci. 2018;38(1):14–25.

11. Cornell-Bell AH, Finkbeiner SM, Cooper MS, Smith SJ. Glutamate induces calcium waves in cultured astrocytes: long-range glial signaling. Science. 1990;247(4941):470–473.

12. Khakh BS, McCarthy KD. Astrocyte calcium signaling: from observations to functions and the challenges therein. Cold Spring Harb Perspect Biol. 2015;7(4):a020404.

13. Shigetomi E, Bushong EA, Haustein MD, et al. Imaging calcium microdomains within entire astrocyte territories and endfeet with GCaMPs expressed using adeno-associated viruses. J Gen Physiol. 2013;141(5):633–647.

14. Araque A, Carmignoto G, Haydon PG, et al. Gliotransmitters travel in time and space. Neuron. 2014;81(4):728–739.

15. Poskanzer KE, Yuste R. Astrocytes regulate cortical state switching in vivo. Proc Natl Acad Sci USA. 2016;113(19):E2675–E2684.

16. Bindocci E, Savtchouk I, Liaudet N, et al. Three-dimensional Ca²⁺ imaging advances understanding of astrocyte biology. Science. 2017;356(6339):eaai8185.

17. Stobart JL, Ferrari KD, Barrett MJP, et al. Cortical circuit activity evokes rapid astrocyte calcium signals on a similar timescale to neurons. Neuron. 2018;98(4):726–735.

18. Semyanov A, Verkhratsky A. Astrocytic processes: from tripartite synapses to the active milieu. Trends Neurosci. 2021;44(10):781–792.

19. Nimmerjahn A, Mukamel EA, Schnitzer MJ. Motor behavior activates Bergmann glial networks. Neuron. 2009;62(3):400–412.

20. Kanemaru K, Sekiya H, Xu M, et al. In vivo visualization of subtle, transient, and local activity of astrocytes using an ultrasensitive Ca²⁺ indicator. Cell Rep. 2014;8(1):311–318.

21. Agarwal A, Wu PH, Hughes EG, et al. Transient opening of the mitochondrial permeability transition pore induces microdomain calcium transients in astrocyte processes. Neuron. 2017;93(3):587–605.

22. Ito M. Cerebellar circuitry as a neuronal machine. Prog Neurobiol. 2006;78(3-5):272–303.

23. Albus JS. A theory of cerebellar function. Math Biosci. 1971;10(1-2):25–61.

24. Eccles JC, Llinás R, Sasaki K. The excitatory synaptic action of climbing fibres on the Purkinje cells of the cerebellum. J Physiol. 1966;182(2):268–296.

25. Marr D. A theory of cerebellar cortex. J Physiol. 1969;202(2):437–470.

26. Palay SL, Chan-Palay V. Cerebellar Cortex: Cytology and Organization. Springer-Verlag; 1974.

27. De Zeeuw CI, Hoebeek FE, Bosman LW, et al. Spatiotemporal firing patterns in the cerebellum. Nat Rev Neurosci. 2011;12(6):327–344.

28. Balakrishnan, S., & Bellamy, T. C. Depression of parallel and climbing fiber transmission to Bergmann glia is input specific and correlates with increased precision of synaptic transmission. Glia. 2009;57(4):393–401.

29. Beierlein M, Regehr WG. Brief bursts of parallel fiber activity trigger calcium signals in Bergmann glia. J Neurosci. 2006;26(26):6958–6967.

30. Cui G, Jun SB, Jin X, et al. Concurrent activation of striatal direct and indirect pathways during action initiation. Nature. 2013;494(7436):238–242.

31. Gunaydin LA, Grosenick L, Finkelstein JC, et al. Natural neural projection dynamics underlying social behavior. Cell. 2014;157(7):1535–1551.

32. Taniguchi H, He M, Wu P, et al. A resource of Cre driver lines for genetic targeting of GABAergic neurons in cerebral cortex. Neuron. 2011;71(6):995–1013.

33. Chen TW, Wardill TJ, Sun Y, et al. Ultrasensitive fluorescent proteins for imaging neuronal activity. Nature. 2013;499(7458):295–300.

34. Aston-Jones G, Cohen JD. An integrative theory of locus coeruleus-norepinephrine function: adaptive gain and optimal performance. Annu Rev Neurosci. 2005;28:403–450.

35. Chandler DJ, Gao WJ, Waterhouse BD. Heterogeneous organization of the locus coeruleus projections to prefrontal and motor cortices. Proc Natl Acad Sci USA. 2014;111(18):6816–6821.

36. Schwarz LA, Miyamichi K, Gao XJ, et al. Viral-genetic tracing of the input-output organization of a central noradrenaline circuit. Nature. 2015;524(7563):88–92.

37. Carey MR, Regehr WG. Noradrenergic control of associative synaptic plasticity by selective modulation of instructive signals. Neuron. 2009;62(1):112–122.

38. Jirenhed DA, Bengtsson F, Hesslow G. Acquisition, extinction, and reacquisition of a cerebellar cortical memory trace. J Neurosci. 2007;27(10):2493–2502.

39. Schonewille M, Gao Z, Boele HJ, et al. Reevaluating the role of LTD in cerebellar motor learning. Neuron. 2011;70(1):43–50.

40. Feng J, Zhang C, Lischinsky JE, et al. A genetically encoded fluorescent sensor for rapid and specific in vivo detection of norepinephrine. Neuron. 2019;102(4):745–761.

41. Yu X, Taylor AMW, Nagai J, et al. Reducing astrocyte calcium signaling in vivo alters striatal microcircuits and causes repetitive behavior. Neuron. 2018;99(6):1170–1187.

42. Strehler EE. Plasma membrane calcium ATPases: from generic Ca2+ sump pumps to versatile systems for fine-tuning cellular Ca2+. Biochem Biophys Res Commun. 2015;460(1):26–33.

43. Ye L, Orynbayev M, Zhu X, et al. Ethanol abolishes vigilance-dependent astroglia network activation in mice by inhibiting norepinephrine release. Nat Commun. 2020;11(1):6157.

44. Ito M. Cerebellar long-term depression: characterization, signal transduction, and functional roles. Physiol Rev. 2001;81(3):1143–1195.

45. Garcia-Garcia MG, Kapoor A, Akinwale O, et al. A cerebellar granule cell-climbing fiber computation to learn to track long time intervals. Neuron. 2024;112(16):2215–2230.

46. Gunaydin LA, Yizhar O, Berndt A, et al. Ultrafast optogenetic control. Nat Neurosci. 2010;13(3):387–392.

47. Lin JY, Lin MZ, Steinbach P, Tsien RY. Characterization of engineered channelrhodopsin variants with improved properties and kinetics. Biophys J. 2009;96(5):1803–1814.

48. Sutton RS, Barto AG. Reinforcement Learning: An Introduction. MIT Press; 2018.

49. Mnih V, Badia AP, Mirza M, et al. Asynchronous methods for deep reinforcement learning. Proc Int Conf Mach Learn. 2016;48:1928–1937.

50. Silver D, Huang A, Maddison CJ, et al. Mastering the game of Go with deep neural networks and tree search. Nature. 2016;529(7587):484–489.

51. Schultz W, Dayan P, Montague PR. A neural substrate of prediction and reward. Science. 1997;275(5306):1593–1599.

52. Schulman J, Wolski F, Dhariwal P, et al. Proximal policy optimization algorithms. arXiv:1707.06347. 2017.

53. Schulman J, Levine S, Abbeel P, et al. Trust region policy optimization. Proc Int Conf Mach Learn. 2015;37:1889–1897.

54. Gong L, Pasqualetti F, Papouin T, Ching SN. Astrocytes as a mechanism for contextually-guided network dynamics and function. PLoS Comput Biol. 2024;20(5):e1012186.

55. Oberheim NA, Takano T, Han X, et al. Uniquely hominid features of adult human astrocytes. J Neurosci. 2009;29(10):3276–3287.

56. Nedergaard M, Ransom B, Goldman SA. New roles for astrocytes: redefining the functional architecture of the brain. Trends Neurosci. 2003;26(10):523–530.

57. Verkhratsky A, Nedergaard M. Physiology of astroglia. Physiol Rev. 2018;98(1):239–389.

58. Khakh BS, Sofroniew MV. Diversity of astrocyte functions and phenotypes in neural circuits. Nat Neurosci. 2015;18(7):942–952.

59. Allen NJ, Lyons DA. Glia as architects of central nervous system formation and function. Science. 2018;362(6411):181–185.

60. Bekar LK, He W, Nedergaard M. Locus coeruleus alpha-adrenergic-mediated activation of cortical astrocytes in vivo. Cereb Cortex. 2008;18(12):2789–2795.

61. Corkrum M, Covelo A, Lines J, et al. Dopamine-evoked synaptic regulation in the nucleus accumbens requires astrocyte activity. Neuron. 2020;105(6):1036–1047.

62. Takata N, Mishima T, Hisatsune C, et al. Astrocyte calcium signaling transforms cholinergic modulation to cortical plasticity in vivo. J Neurosci. 2011;31(49):18155–18165.

63. Picciotto MR, Higley MJ, Mineur YS. Acetylcholine as a neuromodulator: cholinergic signaling shapes nervous system function and behavior. Neuron. 2012;76(1):116–129.

64. Corkrum M, Rothwell PE, Thomas MJ, et al. Opioid-mediated neuron-astrocyte signaling in the nucleus accumbens. Cells. 2019;8(6):586.

65. Bengio Y, Lecun Y, Hinton G. Deep learning for AI. Commun ACM. 2021;64(7):58–65.

66. Hassabis D, Kumaran D, Summerfield C, Botvinick M. Neuroscience-inspired artificial intelligence. Neuron. 2017;95(2):245–258.

67. Lake BM, Ullman TD, Tenenbaum JB, Gershman SJ. Building machines that learn and think like people. Behav Brain Sci. 2017;40:e253.

68. Marblestone AH, Wayne G, Kording KP. Toward an integration of deep learning and neuroscience. Front Comput Neurosci. 2016;10:94.

69. Wang Q, Ding SL, Li Y, et al. The Allen Mouse Brain Common Coordinate Framework: A 3D Reference Atlas. Cell. 2020;181(4):936–953.e20.

70. Guo Fuglstad J, Saldanha P, Paglia J, Whitlock JR. Histological E-data Registration in rodent Brain Spaces. eLife. 2023;12:e83496.

